# Recruitment of Upper-Limb Motoneurons with Epidural Electrical Stimulation of the Primate Cervical Spinal Cord

**DOI:** 10.1101/2020.02.17.952796

**Authors:** Nathan Greiner, Beatrice Barra, Giuseppe Schiavone, Nicholas James, Florian Fallegger, Simon Borgognon, Stéphanie Lacour, Jocelyne Bloch, Grégoire Courtine, Marco Capogrosso

**Author notes:** Correspondence to: Marco Capogrosso, Nathan Greiner.

## Abstract

Epidural electrical stimulation (EES) of lumbosacral sensorimotor circuits improves leg motor control in animals and humans with spinal cord injury (SCI). Upper-limb motor control involves similar circuits, located in the cervical spinal cord, suggesting that EES could also improve arm and hand movements after quadriplegia. However, the ability of cervical EES to selectively modulate specific upper-limb motor nuclei remains unclear. Here, we combined a realistic computational model of EES of the cervical spinal cord with experiments in macaque monkeys to explore the mechanisms of this modulation and characterize the recruitment selectivity of cervical stimulation interfaces. Our results indicate that interfaces with lateral electrodes can target individual posterior roots and achieve selective modulation of arm motoneurons via the direct recruitment of pre-synaptic pathways. Intraoperative recordings suggested similar properties in humans. These results provide a framework for the design of neuro-technologies to improve arm and hand control in humans with quadriplegia.

## INTRODUCTION

Two decades of preclinical and clinical studies have demonstrated that the delivery of EES to the lumbosacral spinal cord can reactivate spinal sensorimotor circuits after SCI^1–9^. Computational and experimental studies conducted in animal models and humans^10–16^ have brought evidence that EES applied over the lumbosacral spinal cord primarily engages large myelinated afferent fibers running in the posterior roots and dorsal columns of the spinal cord. These fibers form synaptic connections with spinal interneurons and motoneurons located in the grey matter, thereby constituting a gateway to the motor circuits controlling leg muscles^17, 18^. For instance, muscle spindle group-Ia fibers provide monosynaptic excitation to motoneurons^19, 20^ and connect to various populations of interneurons located in the intermediate region of the grey matter^21, 22^. Similarly, muscle spindle group-II fibers, force-sensitive group-Ib fibers (from Golgi tendon organs) and low-threshold cutaneous mechanoreceptor fibers all contact specific populations of interneurons, exerting indirect and direct influences onto motoneurons^23, 24^.

Due to their branching morphology^19, 21, 25^, the artificial recruitment of these fibers supplies synaptic inputs to multiple spinal segments. This divergence of excitatory inputs may limit the ability to engage specific muscles with EES, which could be particularly detrimental to applications aiming at restoring arm and hand function. Nonetheless, the distribution of sensory afferents in the posterior roots can be exploited to steer the modulation exerted by EES towards specific motor nuclei. Indeed, selective stimulation of individual posterior roots is known to evoke efferent volleys in the corresponding ventral roots^26^ (*i.e.* same spinal segment). This principle was used to define stimulation protocols that target individual lumbosacral posterior roots independently, with timings corresponding to the natural dynamics of the segmental motoneuronal activity during walking. Such spatiotemporal patterns of targeted EES induced immediate mitigation of lower-limb motor deficits in both animals and humans^3, 8, 9^.

Similarly to the lumbosacral circuits, the segmental organization of sensorimotor circuits controlling upper-limb muscles^27, 28^ suggests that EES of the cervical spinal segments could facilitate the execution of arm and hand function in people with cervical SCI^29^. However, studies on intraspinal microstimulation, which also engages motoneurons via pre-synaptic pathways^30^, have reported low reproducibility and limited specificity of arm muscle recruitment when applied to the cervical spinal cord of monkeys^30–34^, raising questions on the applicability of this technology to the upper-limb. Here, we hypothesized that the distribution of the sensory afferent fibers in the cervical posterior spinal roots can be leveraged to engage specific arm and hand motor nuclei with EES, via the direct recruitment of primary sensory afferents of individual posterior roots.

To test this hypothesis, we implemented a detailed computational model capable of estimating the recruitment of various populations of cervical nerve fibers and neurons in response to single pulses of EES. Numerical simulations suggested that cervical EES primarily recruits large myelinated afferent fibers in the posterior roots and in the dorsal columns, and is unlikely to recruit motor axons directly. The model predicted that the selective recruitment of individual posterior roots (and thus, to a lesser extent, of upper-limb nuclei) was contingent on the precise position of electrodes on the mediolateral and rostrocaudal axes. Electrophysiological experiments in monkeys and humans confirmed that motoneuron recruitment is achieved via synaptic action, thus providing a conceptual framework to design novel EES technologies to improve upper-limb motor control after cervical SCI.

## RESULTS

### Realistic model of the cervical spinal cord

We combined a volume conductor model of the cervical spinal cord with neurophysical models of nerve fibers and neurons to predict the recruitment of fibers and neurons during cervical EES. We used the finite element method to compute tridimensional electric potential distributions in the volume conductor^35^ and we used the estimated distributions to simulate the electrical behavior of neurons and fibers and assess the occurrence of action potentials induced by electrical stimuli^10, 36, 37^.

The volume conductor comprised the following compartments: grey and white matter, spinal roots, dural sac, dura mater, epidural tissue, vertebrae, electrode contacts, electrode insulating paddle, and a wrapping cylinder of conductive material. We developed a geometrical model to describe these compartments and a corresponding software suite to generate numerical instances automatically from user specifications. We determined the geometrical parameters of the cervical spinal cord of macaque monkeys by combining the anatomical analysis of two preserved spines and computed tomography images (see Methods).

Anatomical analysis (**Figure 1b**) revealed consistent cross-section widths across the two analyzed vertebral columns (largest segments were C6 to C8, > 7.5 mm). The length of the spinal segments exhibited a greater degree of variability albeit with a trend of decreasing length from rostral to caudal segments. We found that the trajectories of the posterior and ventral roots from their entrance in the spinal cord to their exit from the spine through the intervertebral foramina were almost perpendicular to the rostro-caudal axis of the spine for spinal segments C2-C3. These trajectories became more and more downward-oriented for the more caudal segments, exiting the spine at markedly more caudal levels than their segment of connection (**Figure 1a**). Consequently, the T1 spinal segment was approximately located at the same rostro-caudal level as the C7 vertebra.

**Figure 1.**
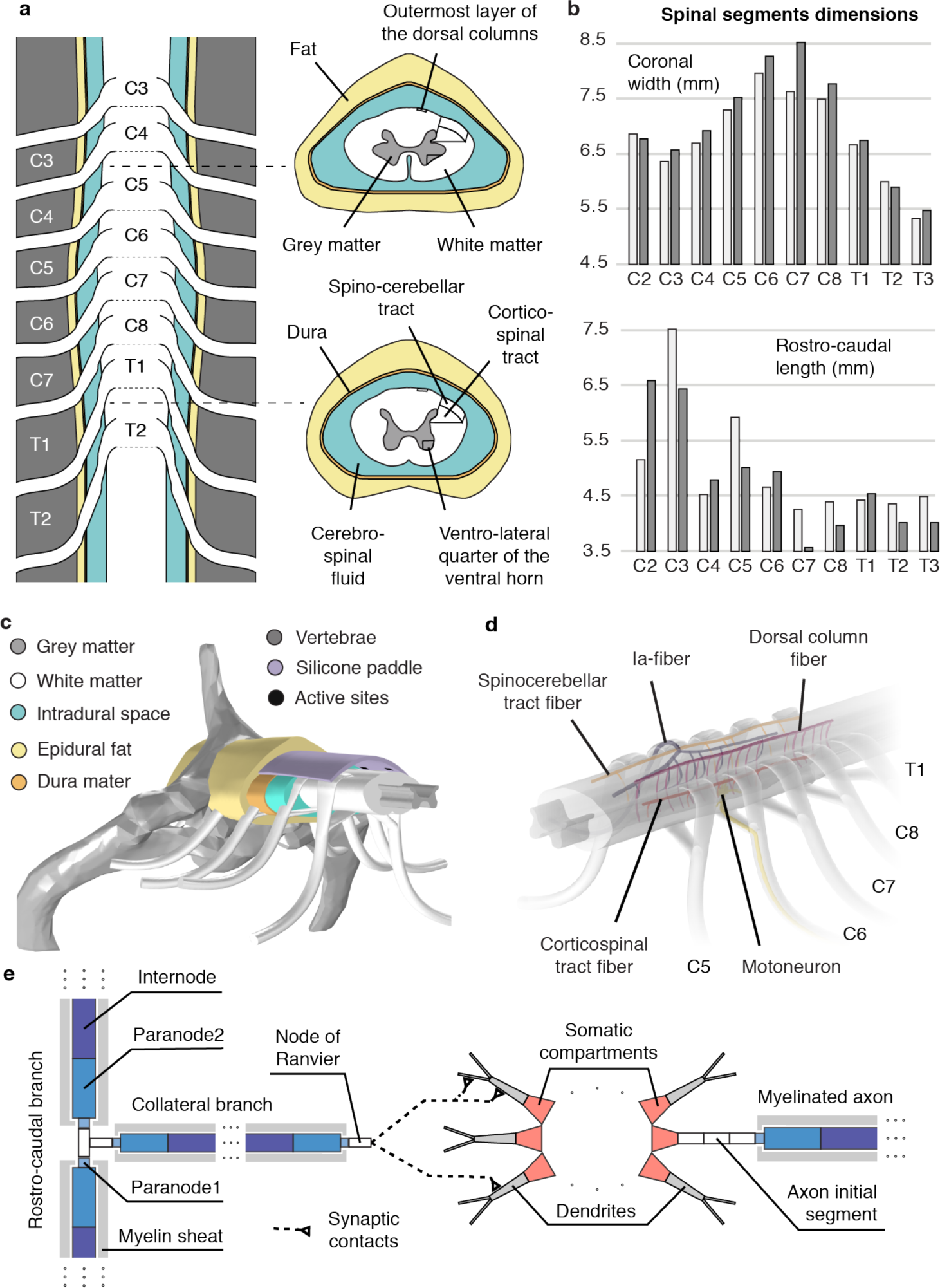
Detailed morphology and computational model of the monkey cervical spinal cord. **a** Macroscopic organization of the cervical spinal cord. *Left:* relationships between spinal segments, spinal roots and vertebrae. *Right:* cross-sections at the C5 and T1 segmental levels showing the internal compartmentalization of the spinal cord. **b** Spinal segments dimensions. The two shades of grey indicate measurements coming from two different spinal cord dissections. **c** Tridimensional view of the volume conductor. **d** Trajectories of virtual nerve fibers and motoneurons. **e** Compartmentalization of myelinated nerve fibers and motoneurons used in neurophysical simulations^32, 37, 39^.

Each modelled tissue was assigned a specific electric conductivity tensor. Importantly, we generated curvilinear coordinates in the white matter and along the posterior and ventral roots in order to appropriately map their anisotropic tensor onto their local longitudinal and transversal directions (see Methods).

We then elaborated realistic tridimensional trajectory models for group-Ia afferent fibers (Aα diameter class), group-II afferent fibers (Aβ diameter class), motoneurons with their efferent axons (Aα), spinocerebellar tract fibers (ST, Aβ), corticospinal tract fibers (CST, Aβ) and dorsal column fibers (DC, Aβ) (**Figure 1c, d**). These were based on documented morphological analyses^21, 25, 38^, and numerical instances were generated at random using stochastic parameters elaborated from these analyses. Motoneurons included dendritic arborizations represented by binary trees of tapered cylinders randomly chosen among a collection of six digital dendritic tree reconstructions. The electrical behavior of nerve fibers and motoneurons in response to extracellular stimulation was emulated with neurophysical compartmental models^32, 37, 39^ using NEURON v7.5 ^40^.

### Computational analysis of the primary targets of EES applied to the cervical spinal cord

We used this simulation environment to estimate the relative recruitment of nerve fibers and cells induced by electrical stimuli delivered from lateral and medial positions of the epidural space.

We hypothesized that 1) laterally-positioned electrodes would primarily recruit large myelinated fibers running in the nearest dorsal roots, 2) medially-located electrodes would primarily recruit dorsal column fibers, and 3) given the dimensions of the primate cervical spinal cord, recruitment of motor axons would not occur within the range of stimulation amplitudes that is necessary to recruit dorsal root afferent fibers.

#### Laterally-positioned electrodes

Our simulations indicate that when placing electrodes laterally, facing a single dorsal root, the strongest generated electrical currents are found directly next to and inside the roots (**Figure 2a**). Furthermore, the generated currents do not substantially penetrate the spinal cord but predominantly flow in the less resistive cerebrospinal fluid. As a result, electric potentials are largest along the dorsal root fibers (DR-fibers, **Figure 2a**), which are recruited at the lowest stimulation amplitudes (**Figure 2b and c**). DR-fibers are followed by longitudinal fibers running in the spinocerebellar tract (ST-fibers), in the dorsal columns (DC-fibers), and in the lateral corticospinal tract (CST-fibers). Finally, the direct recruitment of motoneurons is predicted to remain null even at amplitudes ∼10 times higher than the threshold for Aα-DR-fibers and twice higher than the saturation amplitude for both Aα- and Aβ-DR-fibers and DC-fibers (**Figure 2b**). Surprisingly, the larger diameter of the Aα-DR-fibers does not seem to confer them a substantially lower excitation threshold compared to Aβ-DR-fibers, as could have been expected^41^. This might be due to the close proximity between the electrode and the root, resulting in such steep potential distributions that these diameter differences play a minor role in the fiber excitability.

**Figure 2.**
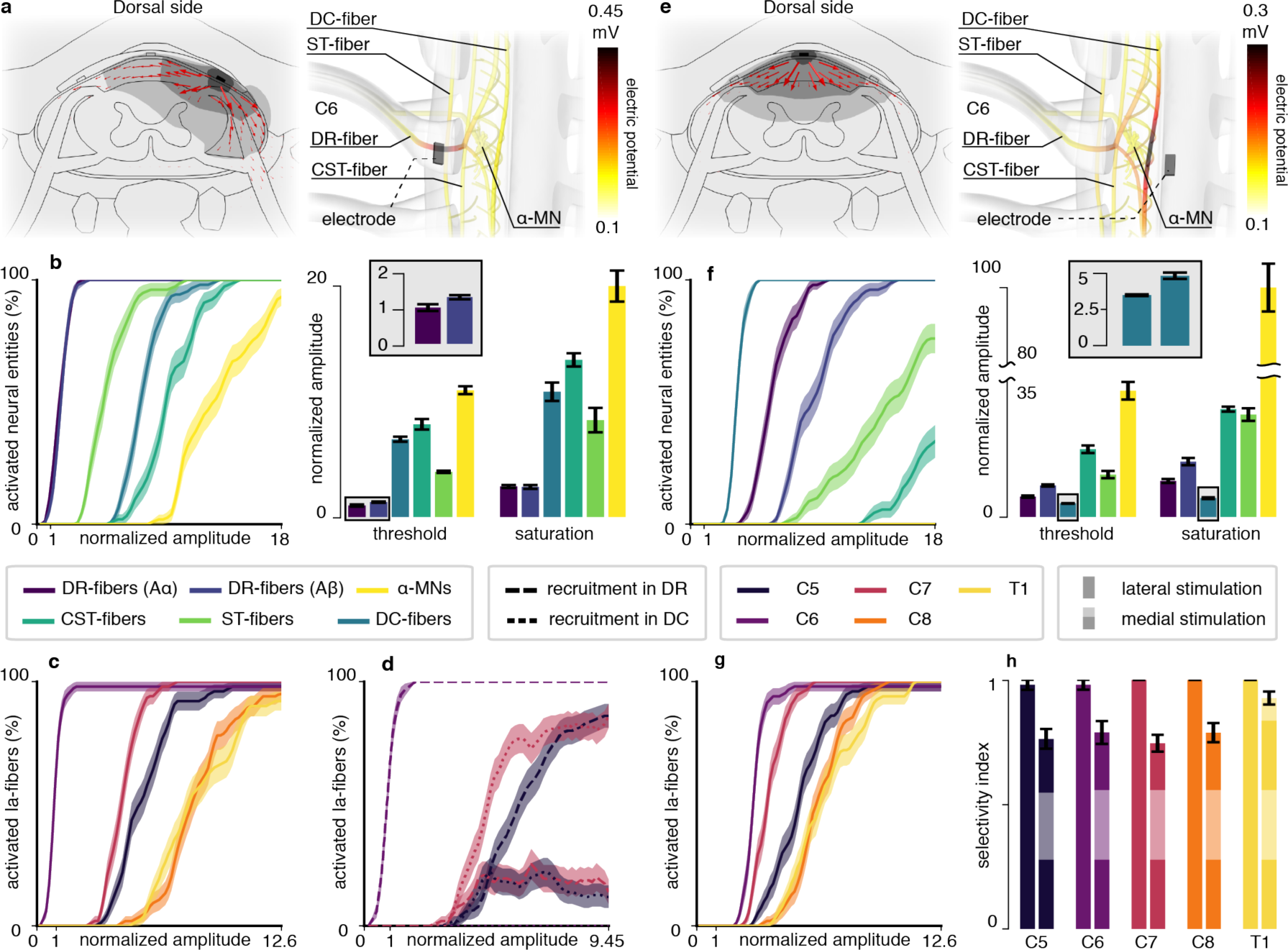
Computational analysis of the direct targets of EES of the monkey cervical spinal cord. **a** Electric currents and potential distribution (ø) generated by a lateral electrode contact at the C6 spinal level for a stimulation current of 1 μA estimated with the finite element method. *Left:* transversal cross-section cutting the electrode contact in two halves. Red arrows: current density vectors. Dark grey surface: ø ≥ 2 mV. Mild grey: ø ≥ 1 mV. Light grey: ø ≥ 0.8 mV. *Right:* ø along trajectories of virtual nerve fibers and motoneurons. DC: dorsal columns. ST: spinocerebellar tract. DR: dorsal root. CST: corticospinal tract. MN: motoneuron. **b** Direct recruitment of nerve fibers and motoneurons estimated from neurophysical simulations using the potential distribution of **a**. *Left:* recruitment curves. *Right:* threshold amplitudes (10% recruitment) and saturation amplitudes (90% recruitment) for the different neural entities, expressed as multiples of the threshold for DR-Aα-fibers. *Inset:* threshold for DR-Aα-fibers and DR-Aβ-fibers. **c** Simulated recruitment of Ia-fibers of individual dorsal roots using the potential distribution of **a**. Amplitudes are expressed as multiples of the threshold for the Ia-fibers of the C6 root. **d** Detailed recruitment for the C5, C6 and C7 roots. Amplitudes are expressed as multiples of the threshold for the Ia-fibers of the C6 root. **e** Same as **a** for a medially-positioned electrode contact. **f** Same as **b** with the potential distribution of **e**. *Inset:* threshold and saturation amplitudes for the DC-fibers. Amplitudes are expressed as in **b**. **g** Same as **c** with the potential distribution of **e**. **h** Maximal selectivity indexes for each root using lateral or medial electrodes (see Methods). *Recruitment curves:* curves are made of 80 data points (except for **d**, 60 data points) consisting in the mean and standard deviation of the recruitment computed across 10,000 bootstrapped populations (see Methods). Lines and filled areas represent the moving average over 3 consecutive data points. *Barplots:* mean and standard deviation across 10,000 bootstrapped populations (see Methods).

We then sought to understand whether DR-fibers of adjacent roots are mostly recruited via the spread of the electric potential towards the adjacent roots or via the recruitment of the dorsal column projections of these fibers.

To this end, we analyzed the recruitment order and action potential initiation sites of Ia-afferents of the C5, C6 and C7 roots in response to stimulation targeting the C6 root.

As expected, our results indicate that Ia-afferents running in the targeted root (C6) are recruited at significantly lower stimulation amplitudes than those running in the adjacent or more distal roots (C5, C7, C8 and T1 **Figure 2c**). Additionally, action potentials are initiated exclusively within their dorsal root branches (**Figure 2d**). In contrast, action potential initiation sites for fibers belonging to the adjacent roots are partly found within their dorsal root branches and partly within their dorsal column projections. Specifically, recruitment of caudal afferents (C7) is mostly induced in their dorsal columns branches, while the situation is reversed for the fibers rostral to the stimulation site (C5). This can be explained by the fact that caudal branches had smaller diameters compared to the rostral branches^19^, increasing their excitation threshold^41^.

#### Medially-positioned electrodes

Electrical currents generated by medial electrodes were predominantly directed towards the dorsal columns (**Figure 2e**), resulting in DC-fibers exhibiting the lowest recruitment threshold (**Figure 2f**), followed by Aα- and Aβ-fibers in the dorsal roots, and finally by ST-fibers. The threshold for CST-fibers is predicted to be ∼5 times that of DC-fibers, while that for motor axons to be as high as ∼9 times this value (**Figure 2f**).

Furthermore, the ability to selectively recruit individual roots was predicted poorer with medial than with lateral electrodes (**Figure 2h**, see Methods). Moreover, this poorer selectivity seems to be mainly due to a less efficient recruitment of the Ia-fibers of the targeted segment (**Figure 2c** and **Figure 2g**).

Finally, analysis of the recruitment order and action potential initiation sites (data not shown) suggested similar recruitment characteristics for caudal and rostral afferents compared to lateral stimulation (**Figure 2d**), although rostral fibers tended to be recruited via their dorsal column projections more often.

Lastly, contrary to lateral contacts, Ia-fibers of the targeted spinal segment were recruited either at the bifurcation point of the dorsal column projections (for most fibers) or the projections themselves.

### Design of cervical spinal implants

Our simulations indicate that, using lateral contacts, it should be possible to recruit the large myelinated fibers of a single root without recruiting fibers of other tracts or roots but marginally. We thus hypothesized that targeting a single root may allow for the selective recruitment of motoneurons innervated by the Ia-fibers of that root. On the other hand, medial stimulation should lead to the activation of a wider range of motoneurons, notably from segments far from the rostro-caudal location of the stimulating electrode.

To obtain experimental evidence supporting this hypothesis, we designed and manufactured two types of multi-electrode spinal implants that were tailored to the dimensions of the spinal cord of macaque monkeys. The first design allowed the delivery of electrical stimuli from both medial and lateral locations of the epidural space. It comprised 2 columns of 5 stimulation active sites: one medial, facing the dorsal columns; and the other lateral, with one active site facing each of the C5 to T1 spinal roots (**Figure 3**). The second design comprised a single, laterally-positioned column of seven stimulation contacts, and one medial stimulation contact spanning three short-circuited active sites (**Figure S1 g and h**)

**Figure 3.**
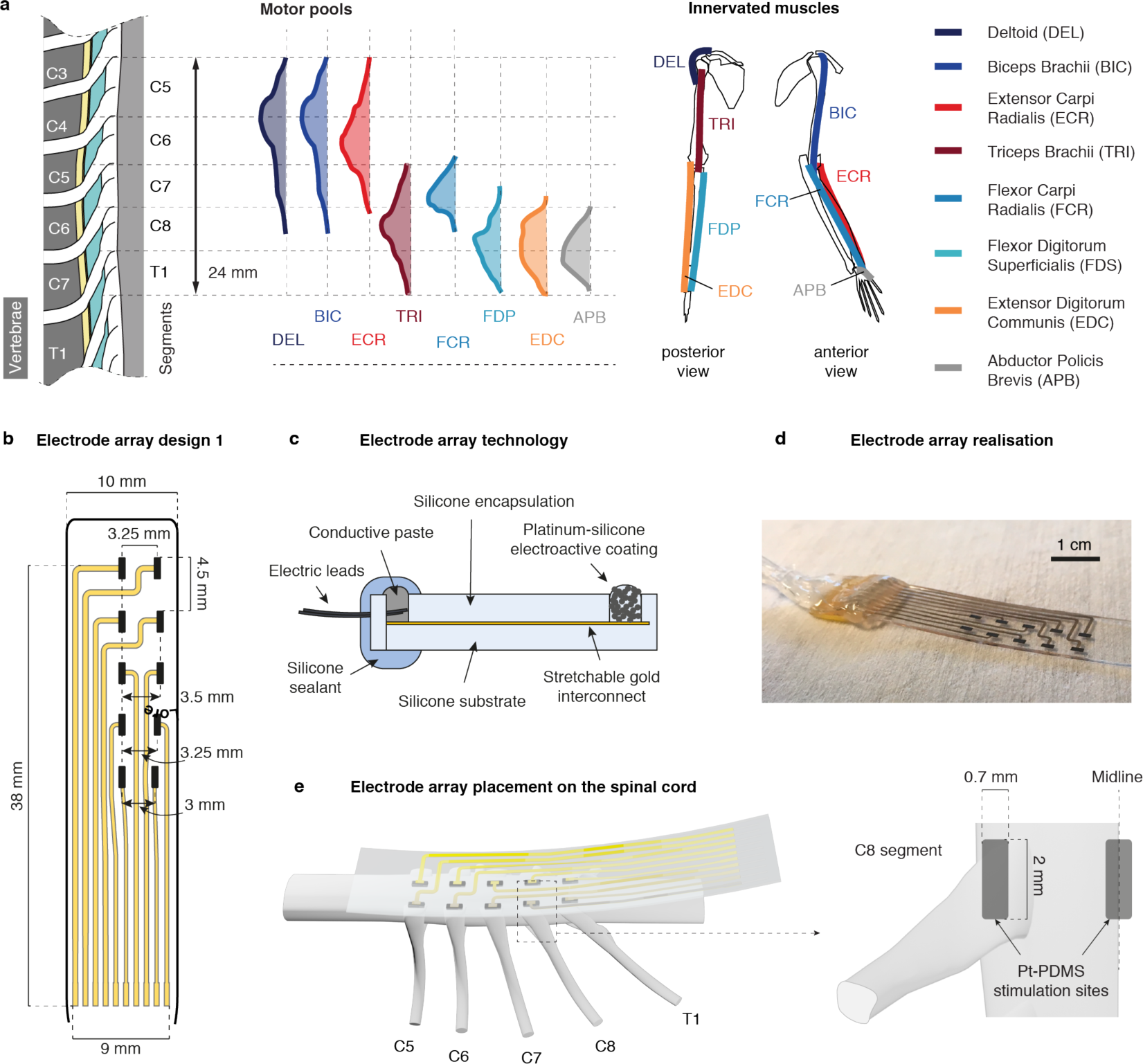
Organization of the monkey cervical spinal cord and soft electrode array tailored to the epidural space of the cervical spinal cord. a. Distribution of the motor pools of 8 upperlimb muscles in the monkey cervical spinal cord^27^ and skeletal positions of these upper-limb muscles. **b** Custom layout of an electrode array with 5 lateral and 5 medial electrode contacts (design 1) tailored to the monkey cervical spinal cord. **c** Cross-section diagram of a soft electrode array with the components labelled. **d** Photograph of a fabricated soft electrode array. *Scale bar:* 1 cm. **e** Placement of the electrode array relative to the cervical spinal cord. Lateral electrode contacts were made to face individual dorsal roots while medial contacts were made to be along the midline of the dorsal columns.

### Experimental recruitment of cervical motoneurons with epidural stimulation in macaque monkeys

#### Laterally-positioned electrodes

We conducted electrophysiological experiments in n=5 anaesthetized monkeys. We implanted the spinal implants (**Figure 3**, n=2 design 1, and n=3 design 2) and delivered single pulses of EES of increasing amplitudes from each lateral contact. In parallel we recorded bipolar electromyographic activity from n=8 muscles of the left arm and hand.

Muscle recruitment curves from monkey Mk-Li (**Figure 4b**) indicate that rostral stimulation (approximately C5 and C6 segments) induced activation predominantly of the deltoid, biceps, and extensor carpi radialis muscles (**Figure 4b**, top panel), all of which are innervated in the C5 and C6 segments. Caudal stimulation (C8-T1 level) mainly recruited the extensor digitorum communis, flexor digitorum profundis, flexor carpi radialis and abductor pollicis brevis muscles (**Figure 4b**, bottom panel), innervated in the C8 and T1 segments (except for the flexor carpi radialis, mostly innervated in C7). Finally, stimulating from an intermediary rostro-caudal level (around C7 level) yielded a recruitment almost purely restricted to the triceps muscle (**Figure 4b**, middle panel), innervated from C7 to T1.

**Figure 4.**
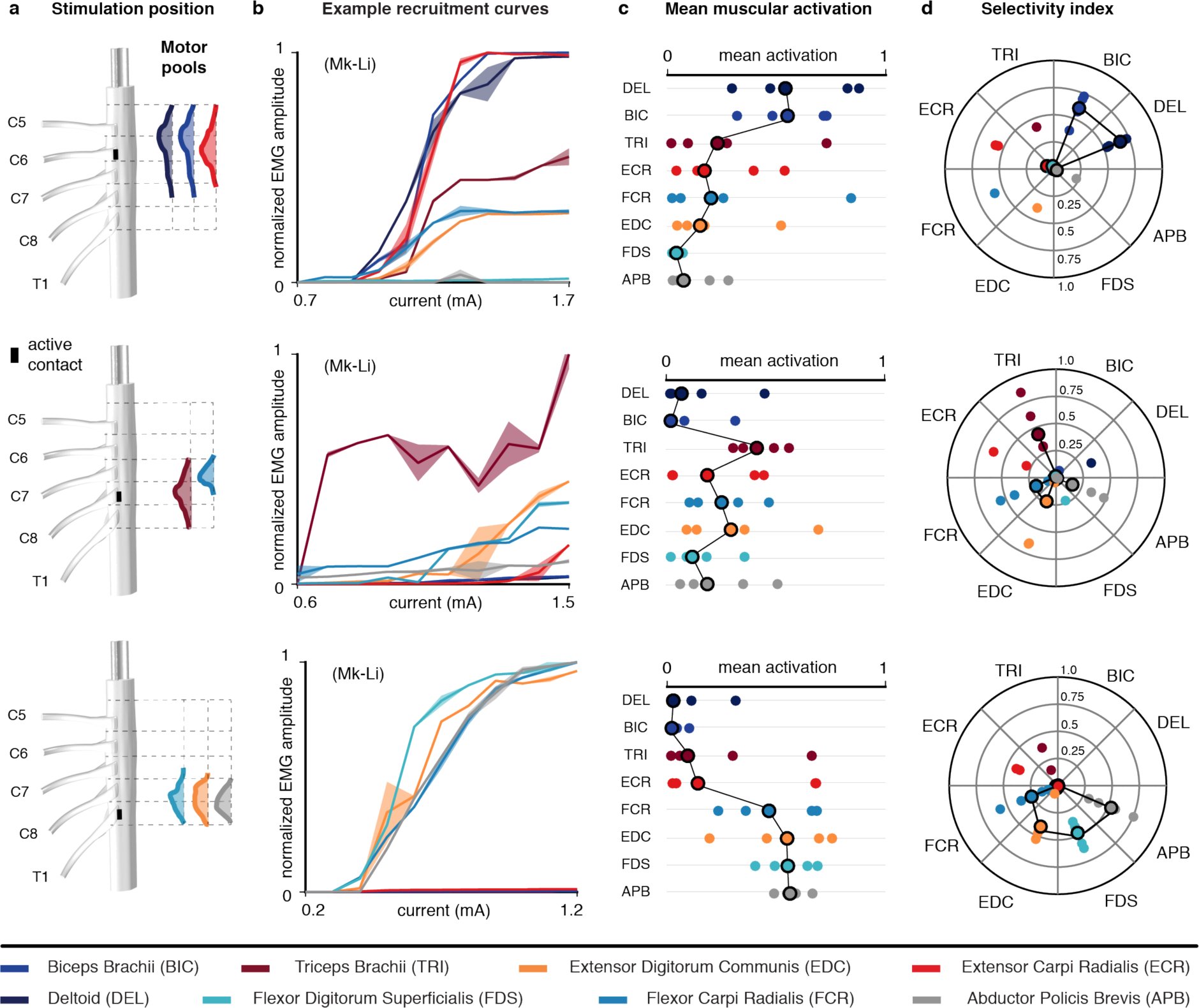
Muscular recruitment induced by laterally-positioned electrode contacts in the cervical spinal cord of monkeys. **a** Approximate positions of the electrode contacts used to obtain the results in **b**, **c** and **d** and underlying motoneuronal distributions. Electrode contacts are magnified for better visualization (scale factor = 2). **b** Examples of muscular recruitment curves observed in monkey Mk-Li using one rostral, one intermediately rostral, and one caudal active contacts. Curves are made of 11 data points consisting of the mean and standard deviation of the normalized peak-to-peak EMG amplitude across 4 responses induced at the same stimulation current. **c** Mean muscular activations observed in 5 monkeys. One rostral, one intermediately rostral and one caudal active contacts were chosen for each animal, and the observed mean muscular activations (see Methods) reported as individual bullets (for Mk-Li, the same active contacts as in **b** were used). **d** Maximal selectivity indexes (see Methods) obtained for each muscle and each animal with the same active contacts as in **c**. *Circled bullets:* medians across the five animals.

Importantly, this rostro-caudal recruitment pattern was obtained with every animal involved in the study, as illustrated by the mean muscular activation levels of **Figure 4c** and the maximal selectivity indexes of **Figure 4d** (see Methods).

#### Medially-positioned electrodes

We then delivered single pulses of EES of increasing amplitudes using medial electrode contacts in n=2 animals (**Figure 5**).

**Figure 5.**
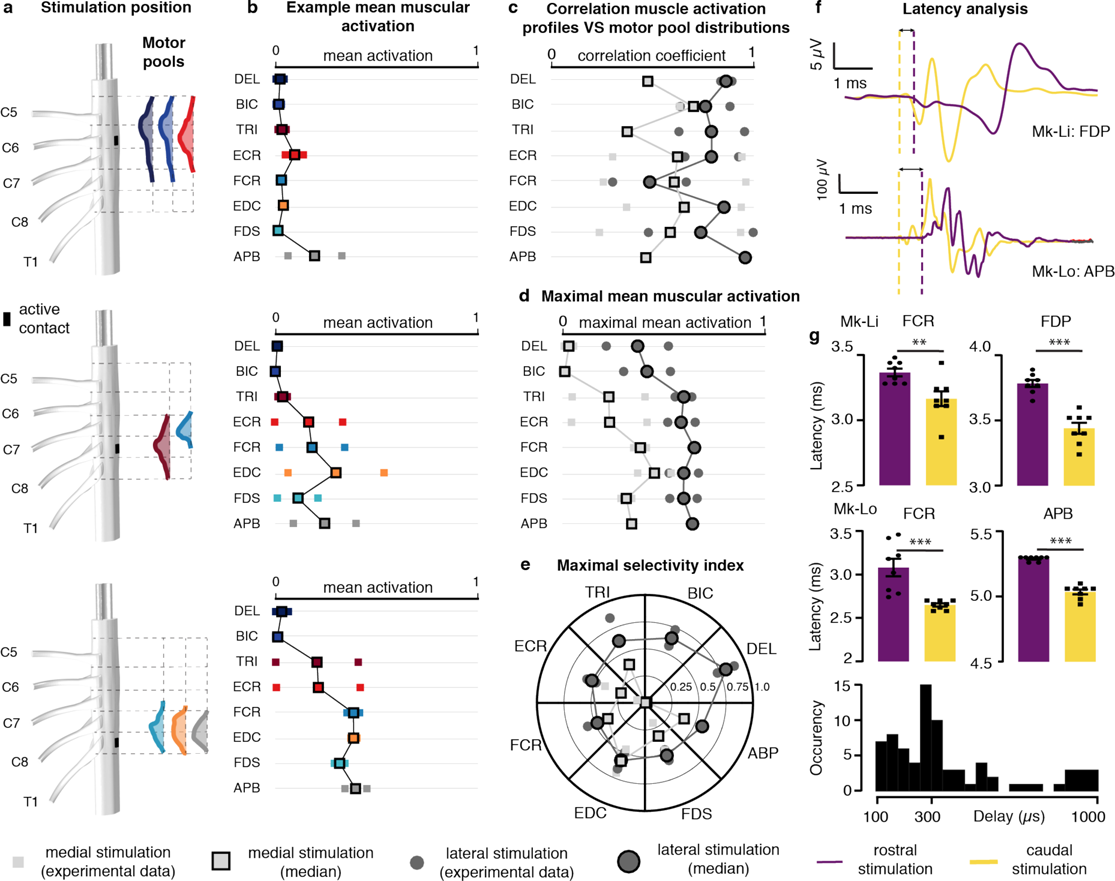
Comparison of muscular recruitment profiles induced by lateral and medial electrode contacts. **a** Approximate positions of the medial electrode contacts used to obtain the results in **b** and underlying motoneuronal distributions. Electrode contacts are magnified for better visualization (scale factor = 2). **b** Mean muscular activations obtained with the medial electrode contacts at the same rostro-caudal levels than the lateral contacts of Figure 4c for the 2 monkeys implanted with the design 1 array (see Methods). **c** Correlation coefficients between muscular recruitment profiles and motor pool distributions (see Methods) for lateral and medial electrode contacts. **d** Maximal mean muscular activations achieved with lateral or with medial electrode contacts. **e** Maximal selectivity indexes (see Methods) achieved with lateral or medial electrode contacts. **f** Examples of muscular response latencies following stimulation from rostral medial or caudal medial electrode contacts. *Top:* muscular responses recorded in the flexor digitorum profundis of monkey Mk-Li. *Bottom:* muscular responses recorded in the abductor policis brevis of monkey Mk-Lo. Onsets of responses are indicated by the vertical dashed lines. **g** Statistical analysis of the differences in onset latencies between rostrally-induced responses and caudally-induced responses. For each muscle, 8 responses induced at amplitudes near motor threshold with one rostral (purple) and one caudal (yellow) active sites were retained. *Bottom histogram:* distribution of the delay of rostrally-induced responses compared to caudally-induced responses.

Compared to lateral, medial stimulation induced a recruitment pattern strongly biased towards caudally-innervated muscles (**Figure 5a,e**). In particular, medial stimulation, even rostral, failed to recruit the deltoid and biceps muscles. Moreover, when computing the correlation between muscle recruitment patterns and motor pool rostro-caudal distributions (**Figure 5d**), lateral stimulation outperformed medial stimulation for all muscles except for the flexor carpi radialis. The maximum level of muscle recruitment (**Figure 5e**) and maximum muscle recruitment specificity (**Figure 5f**) were also higher with lateral compared to medial stimulation.

#### Response Latencies

Finally, we measured the latency of the responses evoked in caudally-innervated muscles following electrical stimulation delivered either from rostral or caudal electrode contacts. We found that rostral stimulation induced responses with latencies that were consistently higher (**Figure 5f,g and S2**). Specifically, the observed differences ranged between 0.2 and 1.1 ms (25^th^ and 75^th^ percentiles respectively) with a median of 0.3 ms. Considering the length separating the rostral and caudal electrodes (approximately 2.5 cm), this means that recruiting a caudally-innervated muscle from a rostral contact elicited responses with an additional latency that matches action potentials propagation velocities ranging from 23 m/s (25^th^ percentile) to 122 m/s (75^th^ percentile) with a median of 71 m/s, which is compatible with diameter-class Aβ-Aα fibers. Furthermore, a symmetrical situation was observed for the biceps muscle in which caudally-induced responses had higher latencies than rostrally-induced responses (consistently across the 2 animals, and with delays comparable with those reported above, **Figure S2**).

### Computational analysis of the Ia-induced motoneuronal recruitment

The previous computational and experimental findings suggested a predominantly Ia-mediated monosynaptic recruitment of motoneurons. We thus sought to use our simulation environment to assess the plausibility of this predominance.

We first assessed the influence of several parameters characterizing the Ia-to-motoneuron synaptic connectivity on the Ia-mediated excitation of motoneurons. These were: the number of Ia-fibers (*n_Ias_*), Ia-to-motoneurons connectivity ratio (*r_connec_*), mean number of synapses per Ia-motoneuron pair (*a_contact_*), and synaptic conductance (*g_syn_*).

Expectedly, we found that the somatic excitatory post-synaptic potential (EPSP) induced in a motoneuron is determined by the product *n_syn_* × *g_syn_* between the total number of synapses it receives (*n_syn_*) and the synaptic conductance (*g_syn_*) (**Figure 6a**). This suggested that the Ia-excitability of a motor nucleus innervated by a population of n_Ias_ Ia-afferents can be characterized by the product *n_Ias_* × *r_connec_* × *a_contact_* × *g_syn_*, since in this case, on average, *n_syn_* = *n_Ias_* × *r_connec_* × *a_contact_* for each motoneuron.

**Figure 6.**
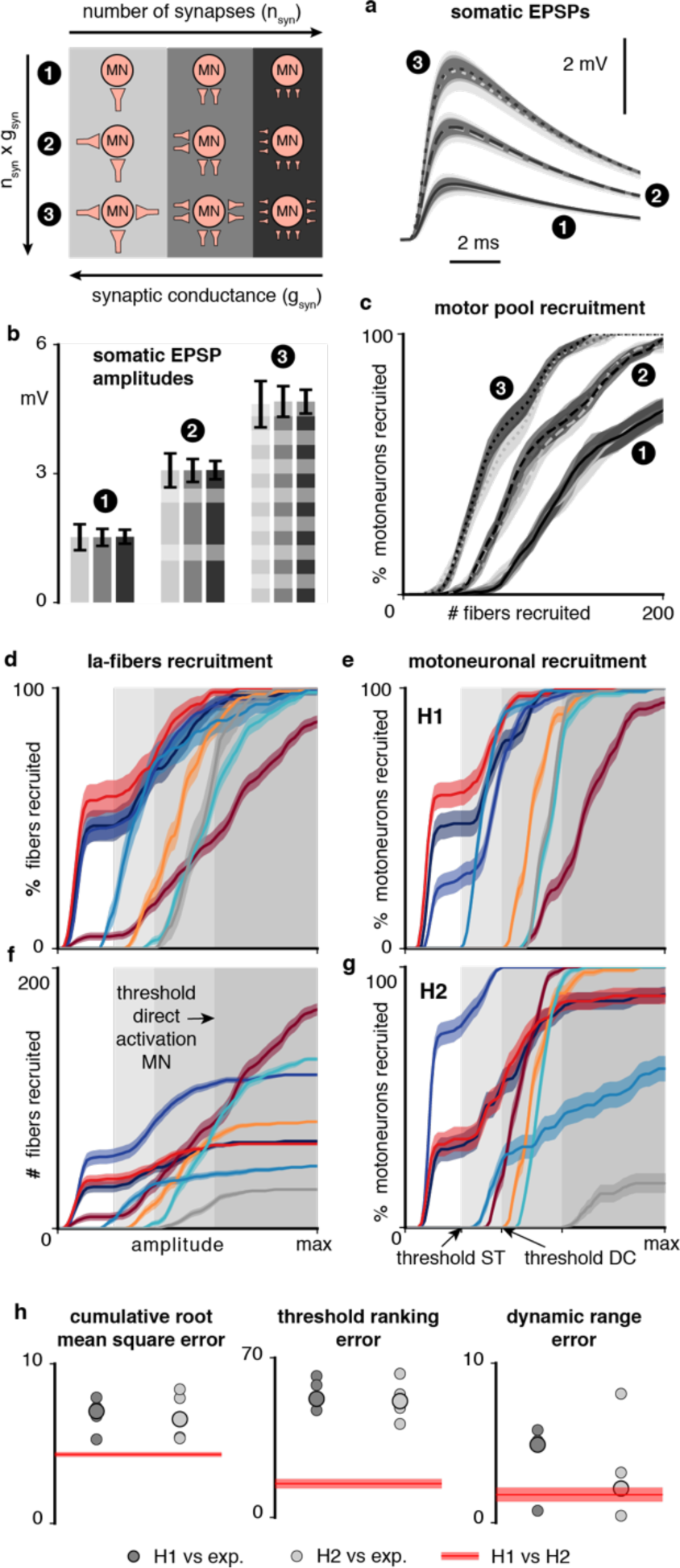
Computational analysis of the Ia-mediated recruitment of motoneurons. **a** Time course of the somatic excitatory post-synaptic potential (EPSP) induced in a motoneuron model by various populations of synapses, for different population sizes *n_syn_* and synaptic conductances *g_syn_*. Lines (solid, dashed and dotted) and filled areas represent the mean and standard deviation of the EPSPs obtained with 100 random synapse populations for each condition (9 conditions in total, see legend and Methods). **b** Maximal amplitudes of the EPSPs of **a**. **c** Recruitment of a motor pool as a function of the number of simultaneously activated fibers innervating this motor pool, for different connectivity ratios and synaptic conductances. Higher connectivity ratios are indicated by higher numbers of synapses in the legend. **d** Recruitment of Ia-fibers of specific muscles following electrical stimulation from a lateral contact at the C6 spinal level (Figure 2a). **e** Monosynaptic recruitment of motoneurons resulting from the Ia-fiber recruitment shown in **d** using muscle-specific synaptic conductances (see **Supplementary Supplementary Table 1**). **f** Same Ia-fiber recruitment as shown in **d** but represented in absolute numbers of recruited fibers. **g** Same as **e** but using a uniform synaptic conductance of 7.625 pS. **h** Comparison between experimental muscular recruitment curves and simulated motoneuronal recruitment curves with H1 or H2 (see Methods). Each bullet represents the comparison score for one animal (5 animals in total). Circled bullets indicate the medians across the 5 animals. *Recruitment curves:* curves are made of 80 data points (except for **c**, 40 data points) consisting in the mean and standard deviation of the recruitment computed across 10,000 bootstrapped populations (see Methods). Lines and filled areas represent the moving average over 3 consecutive data points.

Keeping constant the mean number of synapses per Ia-motoneuron pair (*a_contact_*), we found that the number of Ia-afferents (*n_Ias_*) that needs to be recruited to excite a given motor nucleus is determined by the product *r_connec_* × *g_syn_* (**Figure 6b**). For instance, using *a_contact_* = 9.6 ^19^, *r_connec_* = 0.9 ^25^, and *g_syn_* = 5 pS ^42^, we found that 56 ± 3 (mean ± standard deviation) Ia-fibers were required to induce the recruitment of 10% of their homonymous motoneurons, 93 ± 7 were required to induce the recruitment of 50%, and 175 ± 9 were required to induce the recruitment of 90%.

Next, again assuming a connectivity ratio (*r_connec_*) of 0.9 ^25^ and contact abundance (*a_contact_*) of 9.6 ^19^, and after estimating *n_Ias_* for the 8 upper-limb muscles retained in our study (see Methods), we determined for each of them the minimal *g_syn_* value enabling the recruitment of 100% of its motoneurons. The resulting *g_syn_* values ranged from 3.375 pS (triceps) to 28.5 pS (abductor pollicis brevis) (**Supplementary Table 1**) which are of the same order of magnitude as the 5 pS value estimated experimentally by previous investigators^43^. Thus, our model was able to produce a purely Ia-mediated recruitment of motoneurons within a range of synaptic connectivity parameters coherent with experimental findings.

We then sought to evaluate whether a purely Ia-mediated recruitment is likely to occur during EES. To that end, we evaluated the monosynaptic recruitment of motoneurons resulting from the direct activation of Ia-fibers following lateral stimulation of the C6 root. We either used the *g_syn_* values of **Supplementary Table 1** implying a uniform Ia-excitability across muscles (hypothesis H1), or a uniform average *g_syn_* value for all muscles, implying a higher excitability for muscles possessing higher numbers of Ia-fibers (hypothesis H2) (see Methods). Both hypotheses led to the trans-synaptic recruitment of the motor nuclei located in the targeted segment at stimulation amplitudes that were supra-threshold only for DR-fibers, *i.e.* for group-Ia and group-II fibers (**Figure 6d,f**). Moreover, full (under H1) or almost full (under H2) monosynaptic recruitment could be reached before direct motor axonal recruitment began.

However, comparison between simulated and experimental recruitment curves could not resolve which of the two hypotheses is more realistic (**Figure 6h**).

### Electrophysiological assessment of the trans-synaptic nature of the motoneuronal recruitment

The previous results provide additional support to the hypothesis that the motoneuronal recruitment induced by lateralized EES is predominantly trans-synaptic and Ia-mediated. To gain experimental evidence, we conducted additional electrophysiological assessments. It is well established that repetitive stimulation of primary sensory afferents produces characteristic patterns of responses in the homonymous muscles such as frequency-dependent suppression^44^ or alternation of different reflex responses^45^. We thus delivered supra-threshold stimuli using our spinal implants at multiple locations and frequencies of 10, 20, 50 and 100 Hz in four of our five monkeys. We observed various modulation modalities of the muscular responses in the recorded EMGs as consecutive stimuli were delivered: 1) attenuation of the responses’ amplitude (**Figure 7a**), 2) quasi-suppression of the responses (**Figure 7b**), 3) alternation, often irregular, of two or three stereotypical responses (**Figure 7c**), and 4) erratic adaptation to the pulse trains, wherein the first 5-10 responses didn’t comply to the pattern respected by the subsequent responses of the train (**Figure 7d**). Overall these behaviors were more likely to happen at high frequencies than at low frequencies (**Figure 7f**). Since adaptation of muscular responses should not occur following direct stimulation of motoneurons or motor axons in this range of stimulation frequencies, the previous modulations are all indicative of a mono- or polysynaptic motoneuronal recruitment. Absence of frequency-dependent modulation such as reported in **Figure 7e** was observed only rarely.

**Figure 7.**
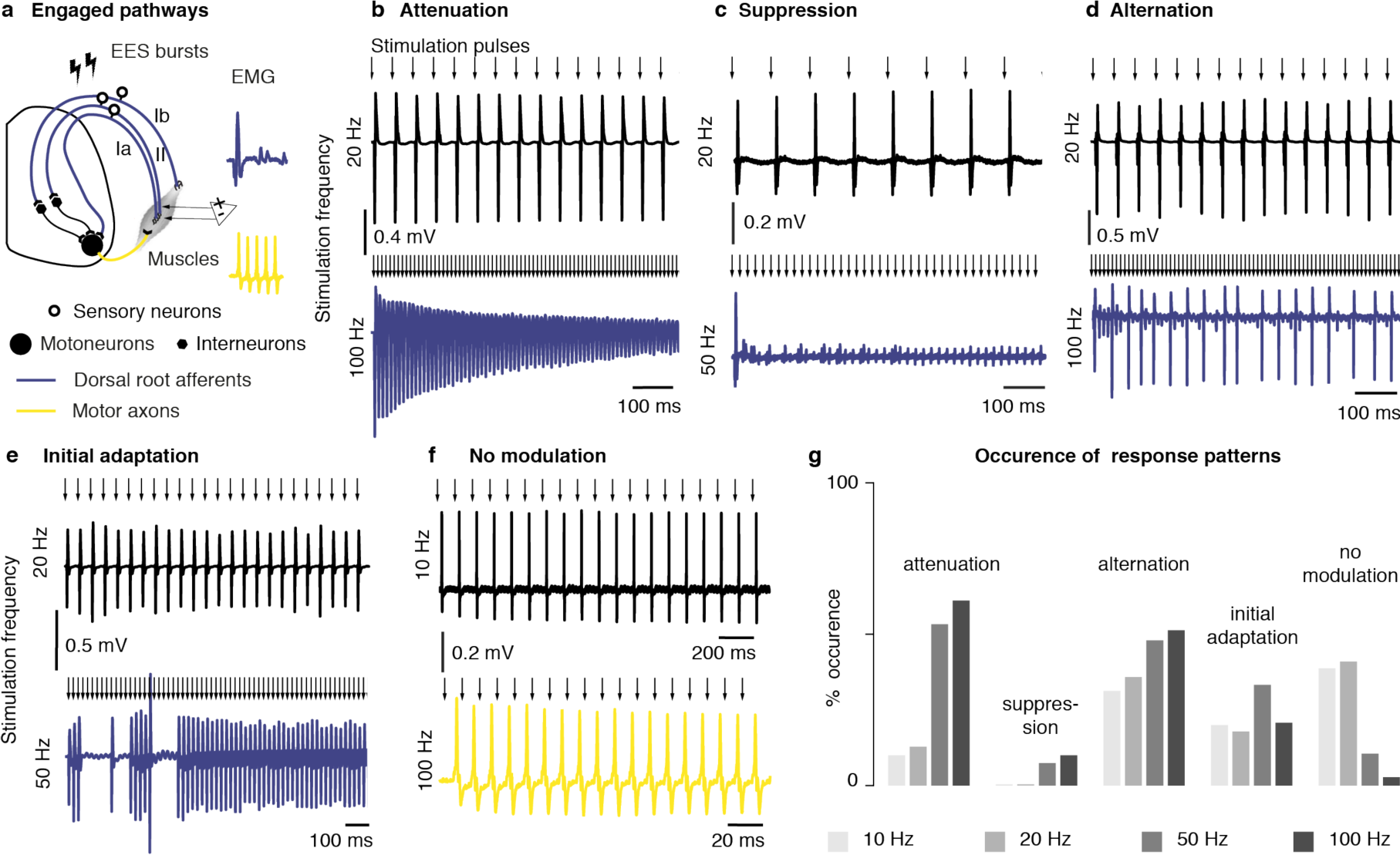
Patterns of muscular responses elicited during high-frequency stimulation of the cervical spinal cord of monkeys. a. Diagram of the presumed engaged pathways during high-frequency stimulation when muscular responses are modulated and unmodulated respectively. **b, c, d, e** Examples of frequency-dependent modulation of muscular responses. In each panel, the top and bottom EMG traces were recorded in the same muscle and using the same stimulation amplitude (near motor threshold) but different frequencies. **f** Example of absence of frequency-dependent modulation. **g** Frequency of occurrence of muscular response patterns during high-frequency stimulation. All the patterns recorded in all the muscles of the 4 animals in which high-frequency stimulation was tested were included in the analysis (n=80 patterns at 10 Hz, n=39 patterns at 20 Hz, n=75 patterns at 50 Hz, n=72 patterns at 100 Hz).

### Analysis of the recruitment of cervical motoneurons with epidural stimulation in humans

We gained access to electrophysiological measurements acquired during clinical procedures involving implantation of a paddle epidural electrode array in three human patients (see Methods). In each patient the paddle array was intended for the treatment of chronic neuropathic pain in the arms and hands, and thus approximately positioned between the spinal segments C6 and T1. Although the clinical procedures did not rigorously match the experimental procedures we performed in monkeys, we could still use the data to conduct some of the analyses we conducted in monkeys. In particular, the clinical tests involved delivering supra-threshold stimuli from individual contacts at frequencies of 1, 10, 20, 60 and 100 Hz. The EMG recordings we analyzed showed muscular responses modulated in a similar way than observed in monkeys. Specifically, we noted: 1) attenuation of the responses’ amplitude 2) alternation, often irregular, of two or three stereotypical responses, and 3) suppression of muscle responses within the first 3-5 responses (**Figure 8c**). Also, as observed in monkeys, modulation occurrences were overall more frequent at high frequencies (**Figure 8d**). In fact, there were no instances of unmodulated responses following stimulation at 60 or 100 Hz. However, we could not look into segmental or muscular specificity since the necessary mapping between maximum muscular contractions and maximum EMG amplitudes was not carried out, preventing the appropriate normalization of the EMG signals and a meaningful analysis of relative muscle recruitment (see Discussion).

**Figure 8.**
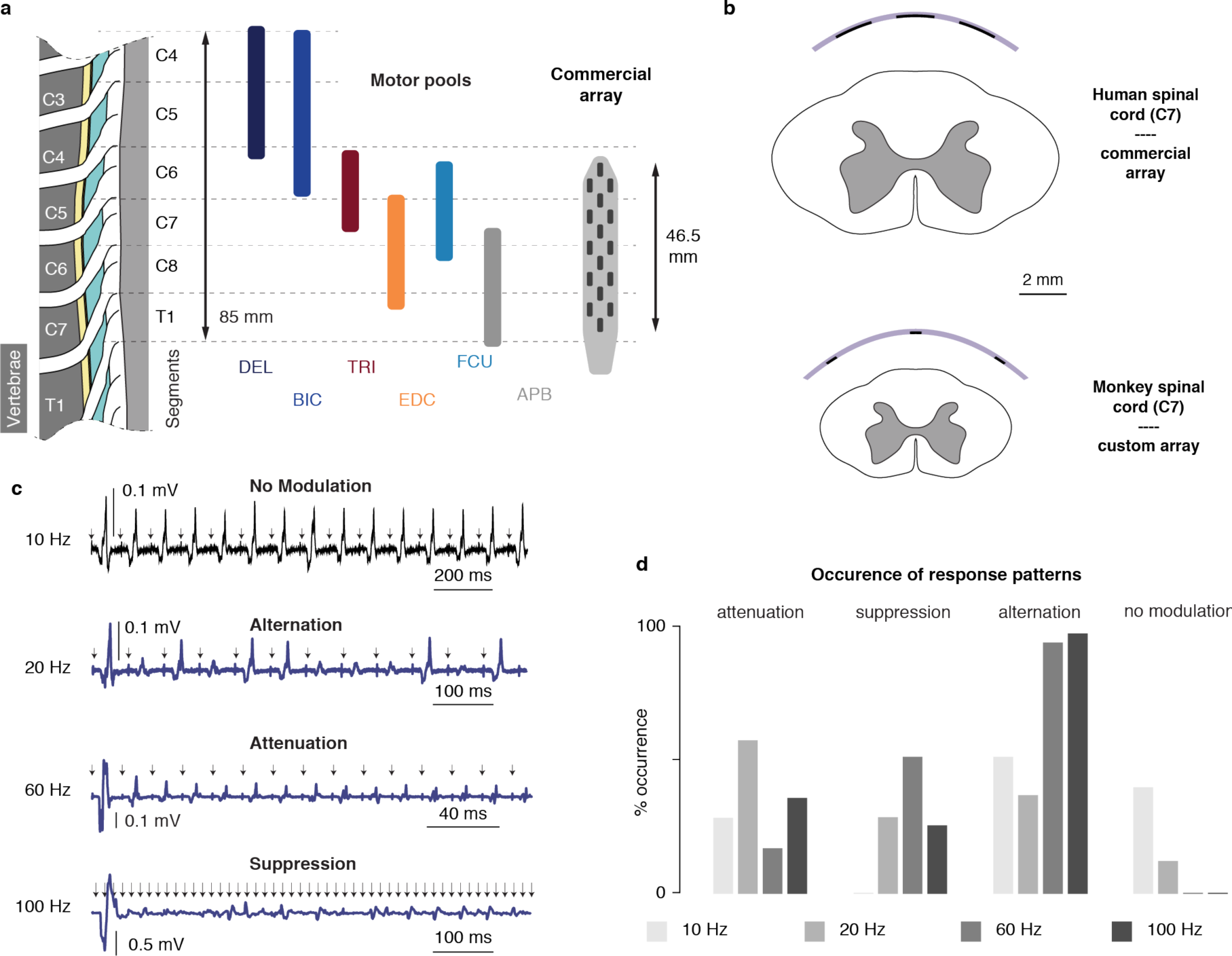
Muscular activity patterns evoked by EES of the cervical spinal cord in humans. **a** Dimensions of the human cervical spinal segments, heuristic distribution of the motor pools of 6 upper-limb muscles^28^, and sketch of the commercial paddle epidural electrode array used in human patients to obtain the data reported in **c** and **d**. **b** Comparison between the relative dimensions of the monkey cervical spinal cord and our custom implant, and the human cervical spinal cord and the commercial epidural implant of **a**. **c** Examples of frequency-dependent modulation of muscular responses. The 4 EMG traces were obtained in the same muscle, using the same stimulation amplitude (near motor threshold) but different frequencies. *Arrows:* timestamps of stimulation pulses. **d** Frequency of occurrence of muscular response patterns during high-frequency stimulation. All the patterns recorded in all the muscles of the 3 subjects were included in the analysis (n=24 patterns at 10 Hz, n=32 patterns at 20 Hz, n=32 patterns at 60 Hz, n=24 patterns at 100 Hz).

## DISCUSSION

In this study, we aimed to identify the main populations of nerve fibers recruited by EES applied to the cervical spinal cord, and to evaluate the influence of the anatomical properties of the cervical spinal cord on the specificity and reproducibility of arm muscle responses that can be achieved with tailored spinal implants.

### Detailed modelling of EES of the cervical spinal cord

The growing development of neuromodulation therapies has accelerated the parallel development and use of computational models to guide the design of neurotechnologies^8, 14, 36, 46, 47^. The recent advent of large computational powers has enabled to increase the realism of the represented neurological systems, which might be critical to obtain accurate quantitative estimates directly usable in clinical applications. For instance, the explicit representation of the spinal roots, which has been missing in most computational models of spinal cord stimulation^10, 13, 14, 16, 36, 48, 49^, should improve the accuracy of the performed simulations. In the present work, we elaborated algorithms to automatically build volumes representing the complex morphology of the roots based on the morphology and dimensions of the spinal cord and vertebral column. We could mesh these volumes with standard commercial software (COMSOL) and generate curvilinear coordinates following their longitudinal course, which allowed the local orientation of an anisotropic conductivity tensor aligned with virtual spinal root fibers. As a result, simulated currents inside the roots were effectively preferentially flowing along the direction of putative fibers. We also devised algorithms to generate realistic trajectories of nerve fibers and cells of various types in the spinal cord and in the spinal roots. In particular, we modelled dorsal root fibers with dorsal column projections and collateral branches projecting towards the grey matter^19, 21, 25, 50^, enabling a finer estimation of the segmental selectivity of the recruitment (see section **Stimulation specificity and lead design**). The computer routines implementing these algorithms are freely distributed and hosted on public online repositories (https://bitbucket.org/ngreiner/fem_smc_ees/src/master/, https://bitbucket.org/ngreiner/biophy_smc_ees/src/master/).

### Trans-synaptic recruitment of motoneurons in the cervical spinal cord

We have found that trains of electrical stimuli (10-100 Hz) modulated the stimulation-induced muscular responses in a frequency-dependent manner, suggesting the trans-synaptic nature of the underlying motoneuronal recruitment. However, in the case of attenuation, alternation, or initial adaptation of muscular responses, the fact that muscle activity is not fully suppressed prevents to completely discard the hypothesis that some direct recruitment of motor axons is occurring. Nonetheless, we believe that our results constitute a strong indicator that the evoked motor responses are predominantly of mono-/poly-synaptic origin, corroborating and extending previous findings^44^.

Our simulation results also suggest that direct motor axonal recruitment is unlikely when using dorsal epidural electrodes. This is attributable to the size of the monkey cervical spinal cord, which leads to relatively low electric potentials in the ventral roots compared to the dorsal roots and the dorsal columns. In turn, this suggests that direct recruitment of motor axons is even less likely in humans, whose spinal cords and vertebral canals are larger while fiber diameters are relatively similar compared to monkeys (thus requiring similar potential distributions for external excitation). At least, our observation that muscular responses induced by cervical EES in humans were also subject to frequency-dependent modulation corroborates the hypothesis of a trans-synaptic recruitment of motoneurons.

### Stimulation specificity and lead design

In our model, the detailed representation of the branching anatomy of dorsal root afferents allowed us to investigate their recruitment via their dorsal column projections. This information is important to determine optimal geometries and placements of epidural electrodes for maximizing the specificity with which muscles can be engaged. Indeed, due to this branching anatomy, our simulations indicate that electrode mediolateral position is a key factor to achieve segmental selectivity (recruiting fibers of individual roots). With medial electrodes, dorsal column fibers are predicted to be more excitable than dorsal root fibers, which is due to the direct exposure of the large cervical dorsal columns to the generated currents^13, 36^. Conversely, laterally-positioned electrodes should be able to recruit all the Ia-afferents of a single root at stimulation amplitudes that are subthreshold for every afferent coming from other roots.

Furthermore, our simulations suggest that the recruitment of afferents of non-targeted roots does not result from the spread of the electric potential along the rostro-caudal direction, but rather from its spread along the mediolateral direction, via the dorsal column projections of these afferents.

Experimental findings were in line with our simulations. The recruitment profiles of arm and hand muscles induced by lateral contacts correlated well with the rostro-caudal distribution of their motor nuclei in the spinal cord. This is in accordance with a selective recruitment of individual dorsal roots under the assumptions that 1) Ia-afferents are distributed in the dorsal roots similarly to their homonymous motoneurons in the spinal segments, and 2) the motoneurons directly innervated by these afferents are predominantly homonymous^9, 10, 19, 20, 51^. Indeed, in this case, if fibers coming from different roots were recruited all at once, stimulation *e.g.* from rostral active sites would not necessarily induce recruitment of rostrally innervated muscles.

By contrast, stimulation from medial contacts elicited markedly distinct muscular recruitment profiles. These profiles correlated less well with the spatial distributions of the motor nuclei, and lower levels of muscular activation were obtained compared to those obtained with lateral contacts. Moreover, the recruitment of caudally-innervated muscles induced by rostral contacts occurred with latencies that were significantly higher than those measured when stimulating with caudal contacts. The differences in latency were compatible with propagation delays through Aβ-Aα fibers, suggesting that rostrally-induced responses in caudally innervated muscles may result from the antidromic recruitment of dorsal column projections of caudal sensory afferents.

The validity of the two assumptions above is not firmly established. First, the distribution of Ia-afferents in the posterior roots in monkeys and humans is largely (and surprisingly) unknown. Second, Ia-afferents are known to also form heteronymous connections^19, 20^. Still, if heteronymous connections remain minority, it seems logical that Ia-afferents enter the spinal cord at the location where their homonymous motoneurons are located. Furthermore, the lack of quantitative information being too important, we did not include heteronymous connections in our model.

Analysis of recruitment selectivity in the clinical data was limited by the nature of the clinical procedures, which were not experimentally-oriented. In particular, we could not reference the muscular activation levels to the maximal EMG amplitudes. Consequently, it was not possible to normalize the recruitment curves of each muscle, and thus to compare the recruitment of different muscles. This limitation prevented us from evaluating the muscular specificity of the stimulation in function of the electrode position. However, comparing the dimensions of the clinical implant and the dimensions of the human cervical spinal cord^52^, it is likely that the electrode array was spanning only few spinal segments and that even the lateral electrodes were facing the dorsal columns. Indeed, existing clinical implants are purposely designed to target dorsal column fibers^49^, suggesting that new implants with lateralized electrodes are required to target individual posterior roots and thus achieve the selective recruitment of motoneurons that we obtained in monkeys with tailored implants.

Despite these limitations, the combination of our simulation and experimental results 1) indicate that epidural stimulation can engage specific arm muscles, 2) are in accordance with a predominantly trans-synaptic recruitment of motoneurons, and 3) suggest that stimulation specificity is limited by the micro-anatomical organization of the dorsal roots. Specifically, the wide spatial separations between adjacent dorsal roots at the cervical level (several millimeters), which is consistent across subjects, should allow robust selective recruitment of individual roots, but the intermingling of different fiber populations within these roots may limit the ability to engage specific motor nuclei.

To further increase the stimulation specificity, a possible strategy would be to target individual rootlets, which might be achieved with high-density electrode arrays and multipolar stimulation configurations. However, the electrical shunting exerted by the dura mater and the cerebrospinal fluid may require to place the electrodes below the dura mater^53^.

### Impact of afferent-to-motoneuron connectivity on muscle recruitments

The trans-synaptic nature of the motoneuronal recruitment during epidural stimulation implies that the connectivity between sensory afferents and motoneurons is of primary importance for the induced muscle recruitment. However, to date, even for the intensively studied Ia-to-motoneuron pathway^19–21, 25, 42, 43, 54^, the properties characterizing the strength of the connection between Ia-afferents and motoneurons remain unknown for most if not all muscles of monkeys and humans.

For instance, while it is well-established that Ia-fibers can induce homonymous motoneuronal recruitment^55^, variability in the Ia-to-motoneuron synaptic connectivity across muscles could influence this recruitment and make some muscles more prone to Ia-mediated activation than others.

Comparison between experimental data and simulation results obtained for different connectivity hypotheses did not allow to discriminate between the tested hypotheses. However, while we simulated the recruitment of motoneurons, the experimental data consisted in normalized EMG peak-to-peak amplitudes, which is only qualitatively related to the total number of recruited motoneurons, a priori disabling a direct comparison. We believe that the lack of thorough and detailed information on afferent-to-motoneuron connectivity currently limits the ability of computer simulations to produce reliable quantitative estimations of EES-induced muscle recruitment and consequently also their use to design personalized clinical therapies.

## Conclusions

By combining computer simulations and electrophysiology in monkeys and humans, we have provided evidence that EES applied to the cervical spinal cord recruits motoneurons trans-synaptically via the direct excitation of sensory afferents. Our results indicate that the selective recruitment of individual posterior roots can be achieved with electrode contacts located on the lateral aspect of the spinal cord. Instead, contacts located over the midline lead to poor segmental specificity, which is due to the recruitment of fibers running in the dorsal columns. These results have direct implications for the design of spinal implants that could be used in human patients to achieve selective recruitment of arm muscles. We believe that these combined results establish a pathway for the development of neuro-technologies for the restoration of arm movements in people with cervical spinal cord injury.

## ACKNOWLEDGEMENTS

The authors would like to acknowledge the financial support from the Wyss Center grant (WCP 008 to GC and MC), the Bertarelli Foundation (Catalyst Fund Grant to MC), an Ambizione Fellowship (No. 167912 to MC), an industrial grant support from GTX Medicals to MC and GC, and the European Union’s Horizon 2020 research and innovation program under the Marie Skłodowska-Curie grant agreement No. 665667 to GS. The authors would like to thank Prof. Eric Rouiller for his support as director of the Platform of Translational Neuroscience of the University of Fribourg; Dr. Eric Schmidlin for kindly providing us with the animals used in the terminal procedures of this manuscript and for his support for the anesthesia; Dr. Melanie Kaeser and Dr. Sara Conti for their assistance during the acquisition of the data in monkeys; Jacques Maillard and Laurent Bossy for their meticulous work on the care provided daily to the animals; and MD Etienne Pralong for providing the clinical data.

## AUTHOR CONTRIBUTIONS

NG and MC conceived the work. NG created and implemented the computational model. NG and SB performed the anatomical analysis. BB, NG, MC, NJ, GC and JB performed the experiments. JB, MC, GC, NG, BB, GS, FF and SL designed the epidural arrays. GS, FF and SL manufactured and tested the epidural arrays. NG, BB and NJ analyzed the data and created the figures. MC and GC supervised the work. NG and MC wrote the manuscript and all the authors contributed to its editing.

## METHODS

### Volume conductor model of the cervical spinal cord

The volume conductor model was implemented in Matlab (Matlab, The Mathworks, Inc.), using COMSOL Multiphysics v5.2a (COMSOL, Burlington MA) to assemble and mesh the geometry, generate the curvilinear coordinates, and compute the electric potential distributions using the finite element method.

Computer code is available at https://bitbucket.org/ngreiner/fem_smc_ees/src/master/.

#### Geometry

Dimensions of the cervical segments were measured from n=2 preserved spinal cords during dissections performed at the University of Fribourg, Switzerland. Preserved vertebral columns were cleared from connective tissue and the spinous processes removed. The dura mater was cut longitudinally and retracted laterally to expose the spinal cord with care to avoid damaging the spinal roots. Spinal roots were labelled according to the vertebral level at which they exited the spine and spinal segments were delimited as portions of the spinal cord extending from the caudal-most rootlet of one root to the caudal-most rootlet of the root directly rostral to it (**Figure 1a**). The segments’ rostro-caudal lengths and coronal widths (at mid-segment-height) were measured independently by 3 experimenters and the average values were retained. We reconstructed cross-sectional contours of the grey matter (GM) and white matter (WM) from a spinal cord atlas^56^ and scaled them using the previous measurements. We formed 3D volumes interpolating these 2D contours using FreeCAD (https://www.freecadweb.org/).

We used OsiriX (Pixmeo SARL) to reconstruct a 3D volume representing the cervical vertebrae obtained from CT-scan images (see imaging section for details). To do so, we first processed the acquired images to correct for the bending of the animal neck during the CT-scan acquisitions using custom Matlab routines. We then used Blender (Blender Foundation) to split the reconstructed volume into individual vertebrae, and MeshLab (Visual Computing Laboratory) to smooth these vertebrae. In this process, their relative positions were preserved.

The GM and WM were positioned with respect to the vertebrae such that the C6 spinal root was perpendicular to the rostro-caudal axis of the spine.

An algorithm similar to ref.^53^ was employed to build the volumes representing the roots. These exited the spine through the intervertebral foramina and had elongated elliptic cross-sections representing rootlets bundles^57^ upon entering the spinal cord.

To build the volumes representing the dural sac, dura mater and epidural tissue, we extracted the vertebrae inner contours which we smoothed and symmetrized using custom Matlab routines. We then scaled them to match the coronal and sagittal widths respectively of the dural sac, dura mater and epidural tissue (determined according to the local spinal cord dimensions), and interpolated them using FreeCAD similarly to the GM and WM. The dura mater thickness was set to 0.15mm ^58^. The dimensions chosen for the dural sac were such that both the dural sac and dura mater were completely enclosed in the vertebral canal and did not overlap with the vertebrae.

The epidural electrode array was represented by metallic active contacts and a large 3D strip embedding the contacts (termed ‘paddle’) in contact with the dura. The dimensions and layout of this array were similar to those of one of the electrode arrays used in the in-vivo experiments presented in this article (design 1). It displayed two parallel columns of 5 active contacts, one along the midline of the spinal cord, and one shifted by ∼3mm on the left side. By design, each row of active contacts was approximately at the rostro-caudal level of one spinal segment among C5 to T1. The active surface of the contacts had a geometric area of 1.0 x 0.5 mm^2^.

Finally, a large cylinder representing the tissues surrounding the spine wrapped all the previous volumes and allowed to apply boundary conditions on its surface ∂Ω.

#### Physics

Each represented tissue was assigned with an electrical conductivity tensor. We used values previously used in literature^53^ for the GM, WM and spinal roots, CSF, epidural tissue, electrode contacts, bone and surrounding saline bath (wrapping cylinder).

We used the Curvilinear Coordinates toolbox of COMSOL to generate curvilinear coordinates in the WM and spinal roots and orient their anisotropic conductivity tensor. Specifically, we used the diffusion method with the inlet set on the top surface of the WM, the outlet set on the surfaces at the tip of each root and the bottom surface of the WM, and the wall set on the remaining lateral surfaces.

Isotropic conductivities of 0.03 S/m ^49^ and 1.0e-13 S/m (typical conductivity for silicone rubbers) were respectively assigned to the dura mater and the electrode paddle.

The capacitive and inductive effects of the materials were neglected, and the quasi-static approximation was employed to compute the electric potential distribution during electrical stimulation^59, 60^. These assumptions lead to condensing Maxwell’s equations into Laplace’s equation 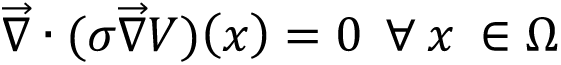 where *V(x)* and *σ(x)* are the electric potential and conductivity tensor at any point x ∈ Ω, and Ω denotes the interior of the volume conductor.

We modeled the delivery of a unitary electric current through the active surface *S* of a contact by imposing 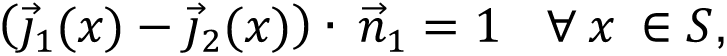 where J⃗_N_, J⃗_P_, and nC⃗_N_ are the current densities on each side of the surface and the normal vector to the surface respectively. The resulting potential distributions (expressed in volts) were then divided by *A*_R_ (the area of S) and by 10^S^, and by considering that they were expressed in millivolts instead of volts, they thus corresponded to a total injected current of 1 μA. We assigned a zero-current flux condition at the outer boundary *δΩ (∇⃗V(x) · n⃗(x) = 0, ∀ x ∈ δΩ)*, and we inserted a grounded point (V_UV2W3X_ = 0) in the ventral region of the wrapping cylinder, playing the role of a virtual return electrode placed there.

This system of differential equations was numerically solved using the finite element method. To this end, the geometry was discretized into a tridimensional mesh of approximately 10 million tetrahedral elements which was denser where high electric potential gradients were expected (near the electrode contacts). The equations’ linearity allowed to estimate the electric potential distributions resulting from arbitrary amounts of injected current as scaled versions of the corresponding unitary distributions.

### Neurophysical models of neural entities

The neurophysical models of nerve fibers and cells were implemented in Python 3.7 (The Python Software Foundation), using NEURON v7.5 ^40^ to solve the membrane potential dynamics. Computer code is available at https://bitbucket.org/ngreiner/biophy_smc_ees/src/master/.

#### α-motoneurons (α-MNs)

α-MNs were modelled with a multi-compartmental soma, a realistic dendritic tree, an explicit axon initial segment, and a myelinated axon. Dendritic trees were derived from digital reconstructions of cat spinal α-MNs established by Culheim and colleagues^61^ freely available from the open-access library NeuroMorpho.org (cell references NMO_00604 to NMO_00609). These specify the geometry of dendritic trees as binary trees of frusta (tapered cylinders) originating at the soma. The axon initial segments comprised 3 linearly-connected identical cylindrical compartments^32^. These were prolonged by a myelinated axon compartmentalized according to the MRG model specifications^62^. The somata were modelled with multiple interconnected frusta following the developments of McIntyre and Grill^32^. For a given α-MN, the soma included one frustum for each dendrite stem, and one for the axon initial segment. The dimensions of the frusta were adjusted to preserve the total area of the soma, imposed by the α-MN diameter. The α-MN diameters were sampled uniformly in the range [44 μm, 71 μm] ^63^. Lengths and diameters of their dendritic compartments were linearly scaled accordingly. Axon diameters were linearly scaled to fall in the range [10 μm, 18 μm]. This last range was an arbitrary estimation for the class Aα fibers in monkeys, which we assumed to be intermediary between that of humans^64^ and that of rats^65^. The dimensions of axon and initial segment compartments were derived from the axon diameters using piecewise linear interpolants established from the data of **Supplementary Table 1** of ref.^37^ and **Supplementary Table 1** of ref.^32^.

The somata centers of α-MNs were uniformly distributed in the ventro-lateral quarter of the GM ventral horn of their host segments. Their dendritic trees, selected uniformly at random among the 6 available templates, were rotated (at random) around their somata centers and partially verticalized to ensure they stayed in the GM or exited it only by short extents. Motor axon trajectories were generated as cubic splines running through the appropriate ventral spinal roots. They kept fixed relative positions along their paths in the spinal roots, reflecting the fact that nerve fibers do not jump from side to sides in nerve bundles.

The biophysical properties of α-MN compartments were the same as in ref.^32^. Numbers of motoneurons and their rostro-caudal distributions for specific motor nuclei were extracted from ref.^27^. For the deltoid (data unavailable) we used the same distribution as for the biceps following observations made in humans^28^.

#### Group-Ia fibers / Dorsal root proprioceptive Aα fibers

Group-Ia fibers were modelled as composed of a dorsal root branch bifurcating into one ascending branch and one descending branch in the dorsal columns^19^ and a series of collateral branches. Collateral branches originated from the dorsal column branches and projected towards the motor nuclei in the GM^21, 25^. Each branch was compartmentalized according to the MRG model specifications^62^. The last node of Ranvier of the dorsal root branch was connected by its extremity to the extremities of the initial nodes of the ascending and descending branches. The initial nodes of the collateral branches were connected to the center of their branching node on the dorsal column branches. The diameters of the dorsal root branches were distributed log-normally with mean μ = 14 μm and standard deviation σ = 3 μm. The diameters of the ascending and descending branches were derived by ponderation by √4/5 and √1/5 respectively. These factors ensured that ascending branches had diameters twice as large as descending branches^19^, and that the total cross-sectional area was preserved upon bifurcation. The number of collaterals *N_cols_* for a given Ia-fiber was estimated based on the data reported by ref.^21^ as *N_cols_* = round(0.75 × L) ± η where L denotes the cumulated rostro-caudal extent of the dorsal columns branches (in mm), η obeys to a Poisson distribution with parameter *λ* = 0.7, and the sign ± denotes an equiprobably positive or negative deviation. Their diameters were log-normally distributed with mean *μ* = 2.5 *μm* and standard deviation σ = 1 μm ^25^. The dimensions of the compartments of the fiber branches were derived from the branch diameters using piecewise linear interpolants established from the data of **Supplementary Table 1** of ref.^37^.

The biophysical properties of Ia-fiber compartments were the same as reported in ref.^37^. In addition, the FLUT compartments contained a fast potassium channel as described in ref.^39^ which was adjusted to a resting potential of −80 mV instead of −70 mV.

Ia-fibers headcounts of the upper-limb muscles were derived from ref.^66^. First, the spindle headcount *sp*^h23^ of muscle *i* was derived from the spindle headcount of the homologous human muscle *sp^hum^_i_ as sp^mom^_i_ = sp^hum^_i_3√m^mon^/m^hum^ where m^mon^ and m^hum^* are typical masses of monkey specimens and human individuals respectively. The group-Ia fiber headcounts were obtained by assuming a 1:1 ratio between Ia-fibers and muscle spindles^64^.

Their distribution in the spinal roots was assumed to be identical to that of their homonymous α-MNs. The resulting distributions, assuming *m^mon^* = 3.5 kg and *m^hum^* = 70 kg, are reported in **Supplementary Table 1**.

#### Dorsal root proprioceptive Aβ fibers

Class Aβ dorsal root proprioceptive fibers were modelled identically to group-Ia / Aα DR fibers, but their diameters were log-normally distributed with mean μ = 9 μm and standard deviation σ = 2 μm.

#### Dorsal columns fibers / Spinocerebellar tract fibers / Corticospinal tract fibers

These fibers were modelled similarly to DR Aβ fibers but they didn’t possess a dorsal root branch. They possessed a rostro-caudal branch running in the dorsal columns, spinocerebellar tract and corticospinal tract respectively. They also possessed a series of collateral branches. The dorsal branches of the dorsal column fibers were restricted to the outermost layer of the dorsal columns, where likelihood of recruitment is highest^67^. Corticospinal tract fibers were meant to represent the large-diameter axons directly connecting to spinal motoneurons and originating from large Layer V cells in the primary motor cortex. Smaller diameter fibers were not represented. Spinocerebellar tract fibers represented Aβ sensory fibers originating from caudal spinal segments.

#### Extracellular stimulation

200 μs-long square pulses of extracellular stimulation were modelled by transiently driving the extracellular batteries of NEURON’s extracellular mechanism to the voltage appropriate for each modelled compartment. The rise and fall of the voltage transients were linear, lasting 2 μs.

#### Synapses

Synapses contacting *α*-MNs were modelled as transient conductances inserted in the membrane of dendritic compartments. The temporal profile of these transients was described by the same function for every synapse, namely *α(t)* = (t - *t_onset_*)_+_ × 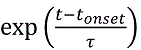 where *t_onset_* is the time onset of the transient (synapse-dependent), τ is the time-to-peak of the transient (synapse-independent), t denotes the running time of the simulation, and (t - *t_onset_*)_+_ is null if t < *t_onset_* and is equal to t - *t_onset_* otherwise. τ was set equal to 0.2 ms ^42^.

Synapses were distributed on the dendritic trees of the contacted α-MNs using the electrotonic-distance-based distribution reported in **Figure 16** of ref.^19^.

The synaptic currents were implemented as *i_syn_*(t) = *g_syn_*α(t)V(t) where *g_syn_* is the maximum of the synaptic conductance transient and V(t) denotes the membrane potential at the synapse location.

For synapses supplied by Ia-fibers, the delay between electrical stimulation and the onset of the conductance transient was the sum of the action potential propagation delay through the Ia-fiber and a log-normally distributed stochastic jitter with mean 1.0 ms and standard deviation 0.5 ms accounting for synaptic transmission. Otherwise, the delay reduced to the previous stochastic jitter.

#### Somatic EPSP analysis

We used a motoneuron model with somatic diameter of 55 μm, 10 dendrites and a total membrane area of ∼475,000 μm^2^. We performed 3 x 3 x 100 simulations where each series of 100 simulations was characterized by a (*n_syn_*, *g_syn_*) pair of values.

These series were grouped by 3, where each group was characterized by a constant *n_syn_* × *g_syn_* product. *Group 1: n_syn_* × *g_syn_* = 500 pS (*n_syn_* = 50, *g_syn_* = 10 pS / *n_syn_* = 100, *g_syn_* = 5 pS / *n_syn_* = 150, *g_syn_* = 3.3 pS). *Group 2: n_syn_* × *g_syn_* = 1000 pS (*n_syn_* = 100, *g_syn_* = 10 pS / *n_syn_* = 200, *g_syn_* = 5 pS / *n_syn_* = 300, *g_syn_* = 3.3 pS). *Group 3: n_syn_* × *g_syn_* = 1500 pS (*n_syn_* = 150, *g_syn_* = 10 pS / *n_syn_* = 300, *g_syn_* = 5 pS/ *n_syn_* = 450, *g_syn_* = 3.3 pS).

#### Ia-mediated monosynaptic recruitment of motoneurons

We assessed the recruitment of a population of 100 motoneurons following the monosynaptic excitation provided by increasing numbers of Ia-fibers (fixed increment of 5 fibers). We assumed a fixed contact abundance of 9.6 synapses per Ia-motoneuron pair, and used 3 x 3 different (*r_connec_*, *g_syn_*) combinations. They were respectively characterized by a product *r_connec_* × *g_syn_* of 3 pS (*r_connec_* = 0.3, *g_syn_* = 10 pS / *r_connec_* = 0.6, *g_syn_* = 5 pS / *r_connec_* = 0.9, *g_syn_* = 3.3 pS), 5 pS (*r_connec_* = 0.3, *g_syn_* = 15 pS / *r_connec_* = 0.6, *g_syn_* = 7.5 pS / *r_connec_* = 0.9, *g_syn_* = 5 pS) or 6.75 pS (*r_connec_* = 0.3, *g_syn_* = 22.5 pS / *r_connec_* = 0.6, *g_syn_* = 12.25 pS / *r_connec_* = 0.9, *g_syn_* = 7.5 pS). The actual number of synapses of a given Ia-motoneuron pair was drawn from a Poisson distribution, and the motoneurons contacted by a given Ia-fiber were drawn uniformly at random.

To evaluate the muscle-specific *g_syn_* values of **Supplementary Table 1,** we assumed a connectivity ratio of 0.9 ^25^, a contact abundance of 9.6 ^19^ and we set *n_Ias_* appropriately for each muscle (see *<u>Group-Ia fibers</u>* section). The *g_syn_* values were determined via a binary search with a resolution of 0.125 pS.

The uniform *g_syn_* value used under H2 was set to 7.625 pS, which is the value estimated for the extensor digitorum communis muscle, possessing an average number of Ia-fibers among the 8 studied muscles.

#### Recruitment curves

Figure 2. Populations of N=50 nerve fibers or motoneurons were simulated for stimulation with multiple electrode contacts and stimulus amplitudes. A nerve fiber or motoneuron was considered recruited when an action potential was elicited and traveled along its entire length.

Threshold and saturation amplitudes of a population were defined as inducing recruitments respectively of 10% and 90% of the population (Figure 2b**,f**). Standard deviations were obtained using a bootstrapping approach (see Statistics section).

Figure 6. Same as above but populations of N=100 motoneurons were simulated for each muscle, and the sizes of the Ia-fiber populations were muscle-specific (see **Supplementary Table 1**).

#### Selectivity indices

The selectivity indices of Figure 2h were computed as

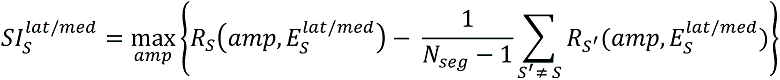

where *R_S_*(amp, E^lat/med^_S_) denotes the recruitment level (between 0 and 1) of the population of Ia-fibers of segment *S* at stimulation amplitude *amp* for either the lateral or medial electrode contact located at the rostro-caudal level of segment *S*.

### Fabrication of custom spinal implants

The implant manufacturing follows the Silicone-on-Silicon process^68^. Implants were prepared on 4” silicon wafers in a class 100 cleanroom environment by using a combination of microfabrication processes adapted to soft materials. The devices are fabricated by processing two identical membranes of silicone elastomer (Polydimethylsiloxane, PDMS, Sylgard 184, Dow Corning), both approximately 220 µm thick. These serve as substrate and encapsulation layers that are subsequently covalently bonded together. A stretchable microcracked thin-film metallization (Cr-Au stack, 5-35 nm) ^69^ is deposited on the substrate by thermal evaporation through a PolyEthylene Terephtalate (PET) mask laminated on the silicone to define the interconnect pattern with conductor tracks leading to different electrodes (**Fig. S1 a**). The encapsulation is laser micromachined to pattern through-holes that serve as vias to access the interconnect at locations corresponding to separate electrodes and wiring pads (**Fig. S1 b**). After the PDMS substrate and encapsulation are processed, they are assembled by covalent bonding following oxygen plasma activation, with the interconnect sandwiched between the two layers and the holes in the encapsulation serving as vias to the exposed gold interconnect (**Fig. S1 c**).

Next, the vias over the electrodes are filled with a soft platinum-PDMS composite material^70^ that provides both high charge injection capacity and mechanical compliance to stretching (**Fig. S1 d**). The implants are then cut to shape and released from the silicon wafer (**Fig. S1 e**). Finally, the vias over the wiring pads are used to make electrical contacts between discrete wires and the gold tracks, using a silver-based conductive paste (Epotek H27D) as soft solder. The connections are then mechanically stabilized by applying a room temperature vulcanization sealant (one component silicone sealant 734, Dow Corning) over the pads (**Fig. S1 f**).

### Electrochemical Impedance Spectroscopy (EIS) of the electrode arrays

EIS measurements were taken *in vitro* by immersing the array under test in a beaker containing Phosphate Buffered Saline solution (Gibco PBS, pH 7.4, 1X), along with a platinum wire as counter electrode and a Ag/AgCl reference electrode (Metrohm, El. Ag/AgCl DJ RN SC: KCl). In this 3-electrode configuration, electrochemical impedance spectra were acquired at room temperature using a Gamry Instruments Reference 600 potentiostat (100 mV amplitude, 1 Hz – 1 MHz frequency).

*In vivo* EIS measurements of implanted electrodes were taken with the same equipment and in the same configuration as *in vitro* measurements. Two separate needle electrodes were inserted percutaneously in the skin on either side of the spine to serve as counter and reference electrodes.

### Experimental procedures

#### Animals involved in the study

Four adult female and one male *Macaca Fascicularis* monkeys were involved in the study. Animal identification and information are summarized in **Supplementary Table 2**. All procedures were carried out in accordance to the Guide for Care and Use of Laboratory Animals^71^ and the principle of the 3Rs. Protocols were approved by local veterinary authorities of the Canton of Fribourg (authorizations reported in **Supplementary Table 2** for each animal) including the ethical assessment by the local (cantonal) Survey Committee on Animal Experimentation and acceptance by the Federal Veterinary Office (BVET, Bern, Switzerland).

Monkeys were housed in collective rooms designed according to European guidelines (45m^3^ for maximum five animals). They had free access to water and were not food deprived. Environmental enrichment was provided in the form of food puzzles, toys, tree branches and devices to climb and hide.

#### Imaging

We performed MRI and CT scans on all animals involved in the study. All procedures were performed at the Cantonal Hospital of Fribourg, Switzerland. The animals were sedated with a combination of ketamine and medetomidine and transported to the imaging facilities where anatomical T1-weighted and T2-weighted images were acquired with a 3T General Electric scanner at a resolution of 0.7 mm. CT scan procedures were analogue to MRI with a spatial resolution of 0.5 mm.

#### Surgical procedures

We performed two types of surgical procedures, terminal in n=3 animals, and acute tests under deep anesthesia in n=2 animals.

Both procedures were performed under full anesthesia induced with midazolam (0.1 mg/kg) and ketamine (10 mg/kg, intramuscular injection) and maintained under continuous intravenous infusion of propofol (5 ml/kg/h) and fentanyl (0.2-1.7 ml/kg/h) using standard aseptic techniques. Animals involved in the terminal procedures were injected with pentobarbital (60 mg/kg) and euthanized at the end of the experiments following the protocols described in the authorizations mentioned in (**Supplementary Table 2**).

During the surgical procedures, the monkeys were implanted with bipolar stainless-steel electrodes to record electromyographic signals (sampling rate = 24 kHz) from the following upper-limb muscles: deltoid (DEL), biceps brachii (BIC), triceps brachii (TRI), flexor carpi radialis (FCR), flexor digitorum superficialis (FDS), extensor carpi radialis (ECR), extensor digitorum communis (EDC), and abductor pollicis brevis (APB). In one monkey (Mk-Ca), the flexor carpi ulnaris (FCU) and extensor carpi ulnaris (ECU) were recorded in place of FCR and ECR respectively. Details on muscle implantation can be found in ref.^72^.

Laminectomies were performed at the T1/T2 and C3/C4 junctions to provide access to the cord and allow insertion of the spinal implants. The custom-made spinal implant was inserted into the epidural space and pulled with the help of a custom-made polyamide inserter. Electrophysiological testing was performed intra-operatively to adjust the position of the electrodes. Specifically, we verified that a single pulse of stimulation delivered through an intermediately rostral electrode induced motor responses in the triceps muscle. Detailed protocols for the implantation and placement of the spinal implant are provided in ref.^72^.

#### Electrophysiology

Trains of biphasic square electrical pulses (charge-balanced, cathodic phase first lasting 200 μs) were delivered at low frequency (0.67 Hz) through a single active site at a time. Within a train, square pulses were grouped by 4 using the same stimulation amplitude, and the amplitude was increased by fixed increments for 11 groups^72^. The first stimulation amplitude was chosen to be the lowest amplitude eliciting a response in any of the recorded muscles, while the last amplitude was chosen by the experimenters upon consensual judgement that some muscle was maximally recruited, either by inspection of the EMG responses or by direct observation of the movements induced in the limbs of the animals.

Additional trains of stimuli were delivered from multiple contacts (at least 2 per animal) at motor threshold stimulation amplitudes and at frequencies ranging from 10 to 100 Hz to test for frequency-dependent modulation of muscular responses^44^.

#### Human data

Anonymized clinical data were obtained from the Centre Hospitalier Universitaire Vaudois (CHUV, Lausanne, Switzerland). The data were obtained during standard clinical practice and under the CHUV’s general ethical approval for clinical procedures and use of data for scientific purposes. The Swiss federal institute of the use of human data for scientific research (Swiss Ethics) validated the use of the anonymized human dataset. Current-controlled electrical stimulation was delivered at frequencies ranging from 1 Hz to 100 Hz and amplitudes ranging from 0 to 5 mA to test for correct surgical placement of Medtronic 5-6-5 lead interfaces in patients with upper-arm neuropathic pain. Percutaneous needles were placed in various upper-limb muscles and EMG signals were recorded (sampling rate = 10 kHz) using a clinical monitoring interface. Exported data were anonymized and provided to the authors with no reference to patient identity.

## Data Analysis

### Muscle recruitment and recruitment curves

From the electromyographic recordings of low-frequency stimulation protocols (0.67 Hz), we extracted 50ms-long snippets of data following each stimulation pulse. For each data snippet, we measured the peak-to-peak amplitude of the recorded signal, *P*2*P*(*s*n*ippe*t) (or 0 V when the signal was purely noisy). Then, for each animal (A), electrode contact (*E*), muscle (*M*) and stimulation amplitude (*amp*), we computed the mean and standard deviation across the 4 *P*2*P*(*s*n*ippe*t) values corresponding to the configuration (A, *E*, *M*, *amp*), respectively noted *P*2*P*^ê^ (*amp*) and *sP*2*P*^ê^ (*amp*). We defined the muscle recruitment *R*^ê^ (*amp*) and associated standard deviation *sR*^ê^ (*amp*) as

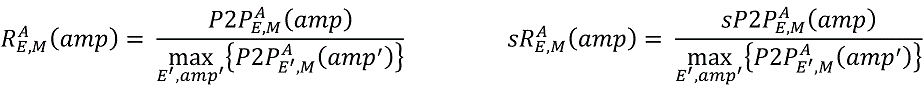

and thereby obtained normalized muscular recruitment curves (as represented in Figure 4b).

### Mean muscle activation and selectivity index

For a given animal (A) and electrode active contact (*E*), the mean activation of muscle *M* was estimated as

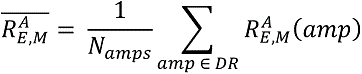

where *DR* denotes the dynamic range of stimulation amplitudes of electrode contact *E* and was defined as the range from threshold amplitude (defined as inducing a muscular recruitment higher than 10% in at least one muscle) to saturation amplitude (defined either as inducing a muscular recruitment higher than 90% in every muscle, or as the highest amplitude used).

The selectivity index of muscle *M* was computed as

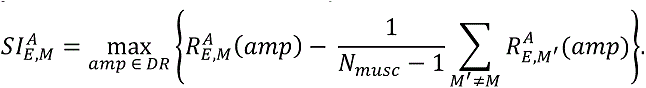

### Comparison between simulated and experimental recruitment curves

*Representative subset of experimental recruitment curves:* for each animal (n=5), we selected 3 recruitment curves respectively obtained with a rostral, a caudal, and an intermediately-rostral lateral active contact.

*Simulated recruitment curves:* the simulated motoneuronal recruitment curves corresponding to the previous 3 active contacts and obtained with each of the two synaptic conductance hypotheses (H1 and H2) were retained for comparison.

*Dynamic range:* here, the saturation amplitude of the simulated and experimental recruitment curves was defined as the amplitude at which any motor pool/muscle was recruited > 90%, incremented by two amplitude steps.

*Cumulative root mean square error (CRMSE):* we restricted the simulated and experimental recruitment curves to their respective dynamic ranges and resampled them over a normalized amplitude vector. The root mean square error (*RMSE*) between the simulated curve with hypothesis *H*, electrode *E* and muscle *M*, and the corresponding experimental curve for animal A was estimated as

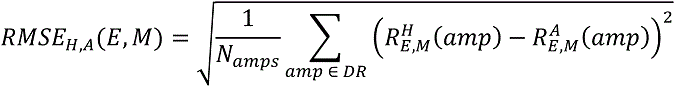

The *CRMSE* between the simulated recruitment curves with hypothesis *H* and the experimental curves of animal A was defined as

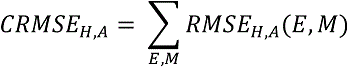

*Threshold ranking error (TRE):* we extracted from the simulated and experimental recruitment curves the recruitment order of the different muscles in the form of lists [*M*_1_, *M*_2_, …, *M_8_]^A/H^_E_* where each *M_i_* denotes a distinct muscle. We then compared the simulated recruitment order under hypothesis *H* with the experimental recruitment order obtained with animal *A* by counting the number of inversions (in the sense of permutations) between the lists [*M*_1_, *M*_2_, …, *M_8_]^H^_E_* and [*M*_1_, *M*_2_, …, *M_8_]^A^_E_*, and summing the obtained values for each of the 3 electrode contact positions.

*Dynamic range error (DRE):* we extracted from the simulated and experimental recruitment curves the normalized length of the dynamic range *L^A/H^_E_* as the length of the absolute dynamic range divided by its first amplitude. The dynamic range error between the simulated recruitment curves under hypothesis *H* and the experimental recruitment curves of animal *A* was computed as

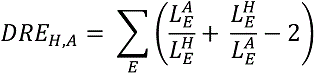

The two sets of simulated recruitment curves were compared with respect to the same 3 metrics. Standard deviations were obtained using a bootstrapping approach (see Statistics section).

### Analysis of frequency-dependent modulation of muscular responses

The recordings of high-frequency (10, 20, 50 and 100 Hz) stimulation protocols from all animals, muscles and electrode contacts were visually inspected and characterized according to 5 criteria: ‘attenuation’, ‘suppression’, ‘alternation’, ‘initial adaptation’ and ‘no modulation’. ‘Attenuation’ and ‘suppression’ applied to those patterns of muscular responses where the amplitude of subsequent responses was progressively reduced and abruptly canceled, respectively (Figure 7a and **b**). ‘Alternation’ applied to patterns where subsequent responses alternated between low amplitude and high amplitude^45^ (Figure 7c). ‘Initial adaptation’ applied to patterns in which the evoked responses became steady only after several (5 – 20) initial pulses (Figure 7d). Finally, ‘no modulation’ applied to patterns in which no observable modulation or change in the evoked responses occurred. Multiple criteria could apply to a single pattern, except for the ‘no modulation’ criteria.

### Analysis of latencies of muscular responses

Latencies of evoked muscular responses were measured as the duration between onset of stimulation and onset of evoked EMG waveform. For every animal and every muscle where this was possible, the difference in latency between the responses evoked by a rostral and a caudal stimulation site was computed (candidate muscles were such that a detectable response could be evoked from both a rostral and a caudal site). In this case, for both the rostral and caudal sites, the latencies of 8 individual responses evoked at stimulation amplitudes near motor threshold were retained to analyze the difference.

### Human data

We re-structured the clinical data provided by the CHUV to analyze it following the same framework used for the monkey data. In particular, we examined recordings of high-frequency (10 to 100 Hz) stimulation protocols to assess the occurrence of frequency-dependent modulation of muscular responses.

## Statistics

### Bootstrapping

For each simulated recruitment curve, the initial population of N simulated nerve fibers or motoneurons was resampled with replacement to obtain K=10,000 fictive populations of N individuals. The means and standard deviations across these fictive populations were used to construct the recruitment curves of Figure 2 and **6**, the threshold and saturation amplitudes and the selectivity indices of Figure 2, and the comparisons between the two sets of simulated recruitment curves of Figure 6h.

