## Supplementary figures and tables for "Recruitment of Upper-Limb Motoneurons with Epidural Electrical Stimulation of the Primate Cervical Spinal Cord"

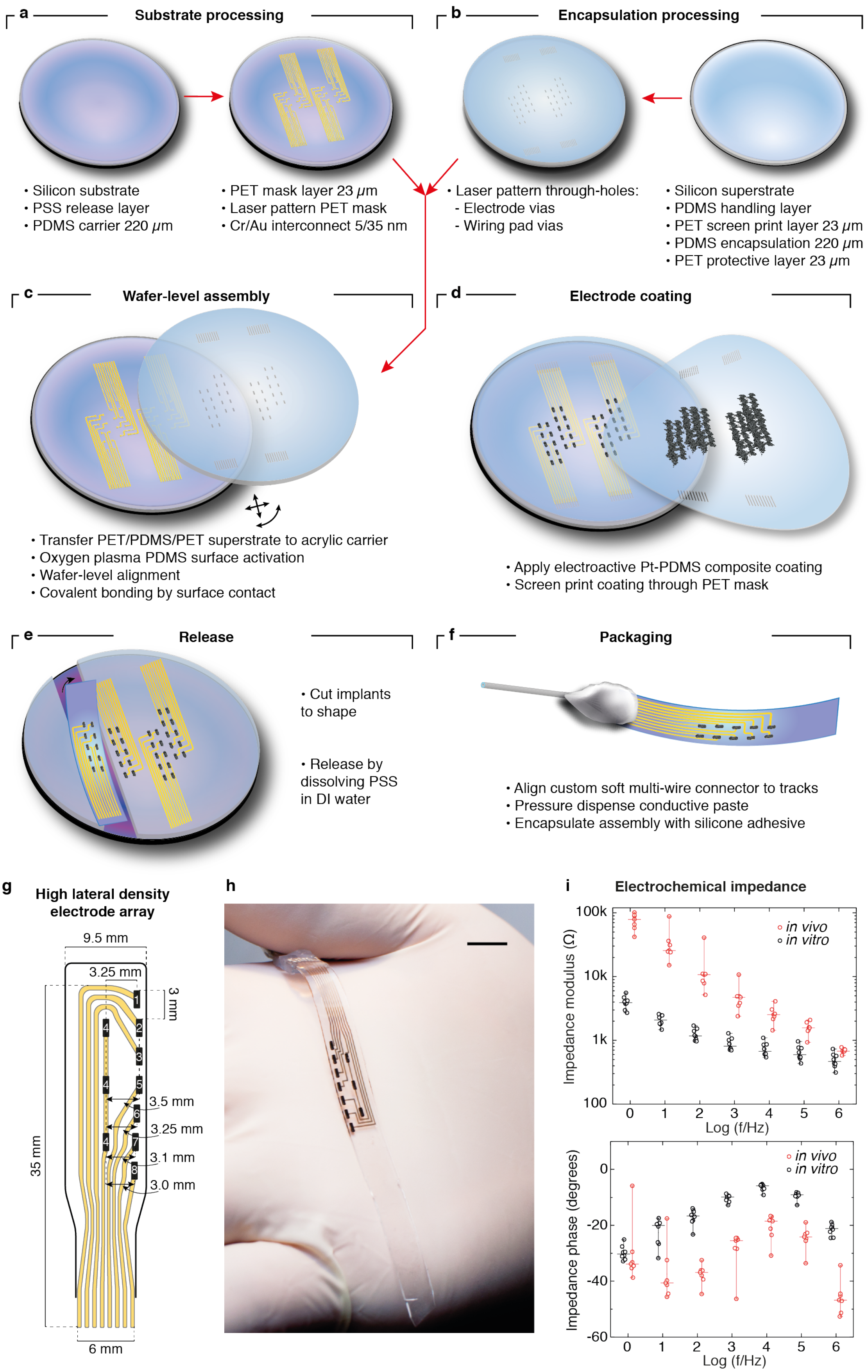
a
Substrate processing
• Silicon substrate
• PSS release layer
• PDMS carrier 220 μm
• PET mask layer 23 μm
• Laser pattern PET mask
• Cr/Au interconnect 5/35 nm
b
Encapsulation processing
• Laser pattern through-holes:
- Electrode vias
- Wiring pad vias
• Silicon superstrate
• PDMS handling layer
• PET screen print layer 23 μm
• PDMS encapsulation 220 μm
• PET protective layer 23 μm
c
Wafer-level assembly
• Transfer PET/PDMS/PET superstrate to acrylic carrier
• Oxygen plasma PDMS surface activation
• Wafer-level alignment
• Covalent bonding by surface contact
d
Electrode coating
• Apply electroactive Pt-PDMS composite coating
• Screen print coating through PET mask
e
Release
• Cut implants to shape
• Release by dissolving PSS in DI water
f
Packaging
• Align custom soft multi-wire connector to tracks
• Pressure dispense conductive paste
• Encapsulate assembly with silicone adhesive
g
High lateral density electrode array
9.5 mm
3.25 mm
3 mm
35 mm
3.5 mm
3.25 mm
3.1 mm
3.0 mm
6 mm
h
i
Electrochemical impedance
Impedance modulus (Ω)
100k
10k
1k
100
Log (f/Hz)
in vivo
in vitro
Impedance phase (degrees)
0
-20
-40
-60
Log (f/Hz)
in vivo
in vitro

**Supplementary Figure 1. Electrode array technology. a-f** Process flow for the fabrication of soft implants on silicon wafers. **a** Substrate processing with the deposition of the stretchable interconnect. **b** Encapsulation processing with the laser micromachining of through-vias. **c** Assembly by covalent bonding of the PDMS substrate and encapsulation. **d** Screen printing of the electrode coating. **e** Implant release from the silicon wafer. **f** Wiring and packaging of the implant. Electrode arrays for cervical neuromodulation. **g** Custom layout of an electrode array with 7 lateral and 1 medial (split in three pieces) electrode contacts (design 2) tailored to the monkey cervical spinal cord. **e** Photograph of a cervical soft electrode array with lateral electrode placement. Scale bar: 1 cm. **f** Electrochemical impedance spectrum acquired on an electrode array implanted in a monkey. Data points are shown at frequencies $f_i = 10^i$, where $i = 0, 1, ..., 6$. Black data points: data *in vitro*, n = 8 electrodes. Red data points: data *in vivo*, n = 7 electrodes. Horizontal bars: means. Whiskers: minima-maxima.

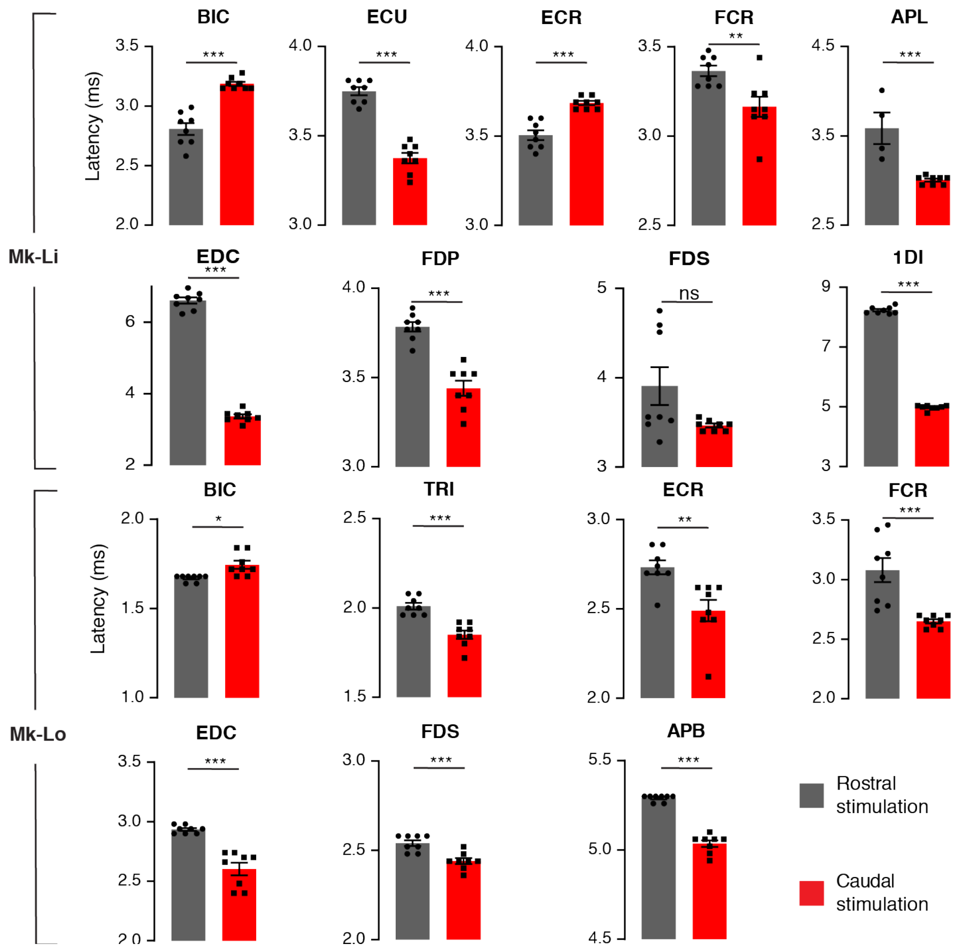


**Supplementary Figure 2. Latency analysis of muscular responses elicited by medial stimulation.** *Top:* latencies of muscular responses recorded in monkey Mk-Li. *Bottom:* latencies of muscular responses recorded in monkey Mk-Lo. For both, all muscles candidate for the analysis were included, and the latencies of 8 responses evoked at stimulation intensities close to motor threshold for one rostral medial and one medial caudal electrode contacts were retained (see Methods). Black-filled circles and squares: data points. Colored rectangles and black bars: means ± SEMs.

**Table 1.** Estimated characteristics of the connectivity between Ia-afferents and motoneurons in 8 upper-limb muscles of the monkey. DEL: deltoid. BIC: biceps. TRI: triceps. EDC: extensor digitorum communis. ECR: extensor carpi radialis. FDS: flexor digitorum superficialis. FCR: flexor carpi radialis. APB: abductor policis brevis.

| Muscle | Ia-fiber headcount | Mean number of synapses per motoneuron | Synaptic conductance [pS] | Total conductance per motoneuron [pS] |
|---|---|---|---|---|
| DEL | 68 | 578 | 9.625 | 5560 |
| BIC | 119 | 1036 | 5.0 | 5180 |
| TRI | 193 | 1681 | 3.375 | 5675 |
| EDC | 82 | 699 | 7.625 | 5330 |
| ECR | 65 | 555 | 10.5 | 5830 |
| FDS | 132 | 1157 | 5.75 | 6650 |
| FCR | 48 | 408 | 15.375 | 6275 |
| APB | 30 | 250 | 28.5 | 7125 |

**Table 2.** Identification information, characteristics, type of procedure performed, and license numbers of the monkeys involved in the study.

| Animal ID | Authorization number | Sex | Age | Weight | Procedure |
|---|---|---|---|---|---|
| Mk-Ca | 2014_42E_ FR | Female | 11 years | 5.8 kg | Terminal |
| Mk-Li | 2017_03_ FR | Male | 12 years | 7.6 kg | Terminal |
| Mk-Cs | 2017_04E_ FR | Female | 9 years | 4.3 kg | Survival |
| Mk-Lo | 2014_42E_ FR | Female | 9 years | 4.0 kg | Terminal |
| Mk-Sa | 2017_04E_ FR | Female | 7 years | 4.4 kg | Survival |
